# A trade-off between stress resistance and tolerance underlies the adaptive response to hydrogen peroxide

**DOI:** 10.1101/2021.04.23.440814

**Authors:** Basile Jacquel, Bor Kavčič, Théo Aspert, Audrey Matifas, Antoine Kuehn, Andrei Zhuralev, Elena Byckov, Bruce Morgan, Thomas Julou, Gilles Charvin

## Abstract

The physiological adaptation to environmental stress involves complex molecular responses leading to separate cellular fates aimed at maximizing fitness: either cells can maintain proliferation by degrading the effects of the stressor (i.e. resistance), or they focus on ensuring cell survival (i.e. tolerance), even at the expense of proliferation. These strategies are complementary, yet whether they are coordinated to ensure an optimal physiological stress response remains unknown. Here, we used microfluidics and live cell imaging to explore the genetic basis of the interplay between resistance and tolerance during the response to hydrogen peroxide (H_2_O_2_) in budding yeast. Our analysis unraveled that the deletion of *zwf1*Δ, which is responsible for NADPH synthesis via the PPP pathway, led to a decrease in resistance that was counterbalanced by an unexpected exacerbation of tolerance to H_2_O_2_. This trade-off between stress resistance and stress tolerance was further characterized using both genetic and environmental interventions, and we confirmed that it was conserved in bacteria. Our results support a model in which redox signaling triggers the switch to a nutrients-dependent non-proliferative tolerant state via inhibition of protein kinase A when the H_2_O_2_ homeostatic response is overwhelmed. Our framework could help develop synergistic therapies that target mechanisms driving both resistance and tolerance to prevent drug escape mechanisms and disease relapse.

## Introduction

Cell responses to stress are complex and involve a variety of mechanisms that promote physiological adaptation in changing environments. A broad class of defense mechanisms against both environmental and endogenous insults uses sense-and-respond regulatory systems based on the homeostatic framework. Many such mechanisms have been described with exquisite molecular detail in budding yeast, such as the oxidative stress response^1–3^ and the response to hyperosmolarity^4^. The role of the homeostatic system is thus to ensure the process of stress adaptation, i.e. to enable cells to recover their canonical physiological functions such as growth and proliferation despite the presence of internal or external disturbances^5,6^.

However, whereas such regulatory mechanisms may drive excellent adaptive properties at steady-state^7–9^, they often suffer from several limitations. These include a limited homeostatic range, a slow response time that may hinder the cell’s ability to cope with abrupt environmental changes^7,10^ and imperfect homeostatic regulation. Consequently, under stress, cells often operate in a state akin to but distinct from homeostasis, known as allostasis^11^. To compensate for the aforementioned limitations of homeostatic systems, stress response encompasses broad transcriptional changes, metabolic rerouting^12,13^, and growth-regulating processes^14,15^ that do not directly participate in stress detoxification. For example, this includes protective mechanisms (such as heat-shock aggregates, stress bodies, etc.) and repair mechanisms (including DNA repair, chaperone proteins, etc.) that mitigate the damages caused by stress.

In microbiology, the physiological adaptation to stress has been conceptualized by distinguishing two distinct properties: stress resistance and tolerance. Resistance refers to the ability of cells to maintain or restore proliferation during continuous stress exposure. Conversely, stress tolerance is defined as the ability of cells to survive a transient physiological threat without necessarily showing a *bona fide* adaptation to the stressor. Both of these phenomenological properties can be independently measured^16^, illustrating distinct defense strategies: cellular resistance may *a priori* maximize cellular fitness, yet it exposes the cells to damage. Instead, tolerance mechanisms, which are usually accompanied by arrested proliferation and reduced metabolism^17^, or heterogeneous cellular behavior such as bacterial persistence^18,19^ and bet-hedging^20,21^, come at the cost of reduced cellular proliferation and offer only temporary protection against stress exposure. This distinction has proven particularly powerful in characterizing cellular strategies in the context of antibiotic exposure but can be extended to any stress response context. However, it remains to understand whether and how each component of the cellular response (e.g. homeostatic system, protection and repair mechanisms, etc) maps onto resistance, tolerance, or both.

Redox homeostasis is an essential feature for cells to function properly when facing redox perturbations of external and internal origin^1,2^. In yeast, the Yap1 regulon controls hydrogen peroxide (H_2_O_2_) levels through a canonical sense-and-respond system^22–24^: upon H_2_O_2_ exposure, the nuclear-sequestered Yap1 transcription factor drives the expression of about one hundred genes ^22,25,26^, including antioxidant enzymes with somewhat overlapping H_2_O_2_ scavenging functions ^27–31^. Additional regulations contribute to the restoration of internal H_2_O_2_ balance: first, glycolysis rerouting to the pentose phosphate pathway (PPP) leads to increased production of NADPH ^32,33^, which is the ultimate electron donor involved in H_2_O_2_ buffering in the peroxidatic cycle^34^. Additionally, the inhibition of the protein kinase A (PKA) pathway, which is a major hub for cell proliferation control^15,35^ and the general stress response^26,36^, contributes to the adaptation to oxidative stress^37^ and is connected to the H_2_O_2_ signaling response through various putative mechanisms^38–41^. Therefore, the response to H_2_O_2_ in yeast provides an ideal context for studying how resistance and tolerance mechanisms shape a multi-faceted stress response.

To address this question, we used live-cell imaging and microfluidics approaches to develop combined proliferation and survival assays that can distinguish between H_2_O_2_-resistant and tolerant cellular behaviors. Using a candidate-gene screen, we classified the main players involved in H_2_O_2_ stress response into functional categories that highlight their respective roles in adaptation to this stressor. Specifically, our study revealed the existence of a trade-off between resistance and tolerance to hydrogen peroxide, which was exacerbated by mutations that affect NADPH fueling in the peroxidatic cycle in budding yeast (e.g., *zwf1*Δ and *trr1*Δ mutants), a phenomenon that appeared to be conserved in *E. coli* when mutating the zwf gene. Further analyses unraveled the role of PKA regulation and the availability of nutrients in orchestrating the interplay between resistance and tolerance upon exposure to H_2_O_2_. This model system paves the way for developing anti-proliferative strategies in which both resistance and tolerance mechanisms could be independently targeted to improve therapeutic efficiency.

## Results

### Resistance and tolerance are distinct physiological properties of the hydrogen peroxide stress response

Classical oxidative stress response assays, typically performed on agar plates or in liquid cultures, often fail to distinguish between a cell’s survival and its capacity to proliferate under stress. In contrast, single-cell assays offer a valuable approach to independently measure proliferation and tolerance, thereby allowing us to determine whether these two properties are truly distinct and, if so, under which specific stress conditions this distinction occurs.

Hence, we developed two independent microfluidics-based single-cell time-lapse assays. In the first, we monitored the proliferation of individual cells and their progeny under a constant concentration of H_2_O_2_ (i.e., resistance assay; see Figure 1A). We then measured the fraction of resistant cells across increasing H_2_O_2_ concentrations (Figure 1B and 1C). This assay revealed a sharp decline in cell proliferation above ∼0.7 mM, a threshold referred to as the minimum inhibitory concentration (MIC; see Figure 1B).

**Figure 1:**
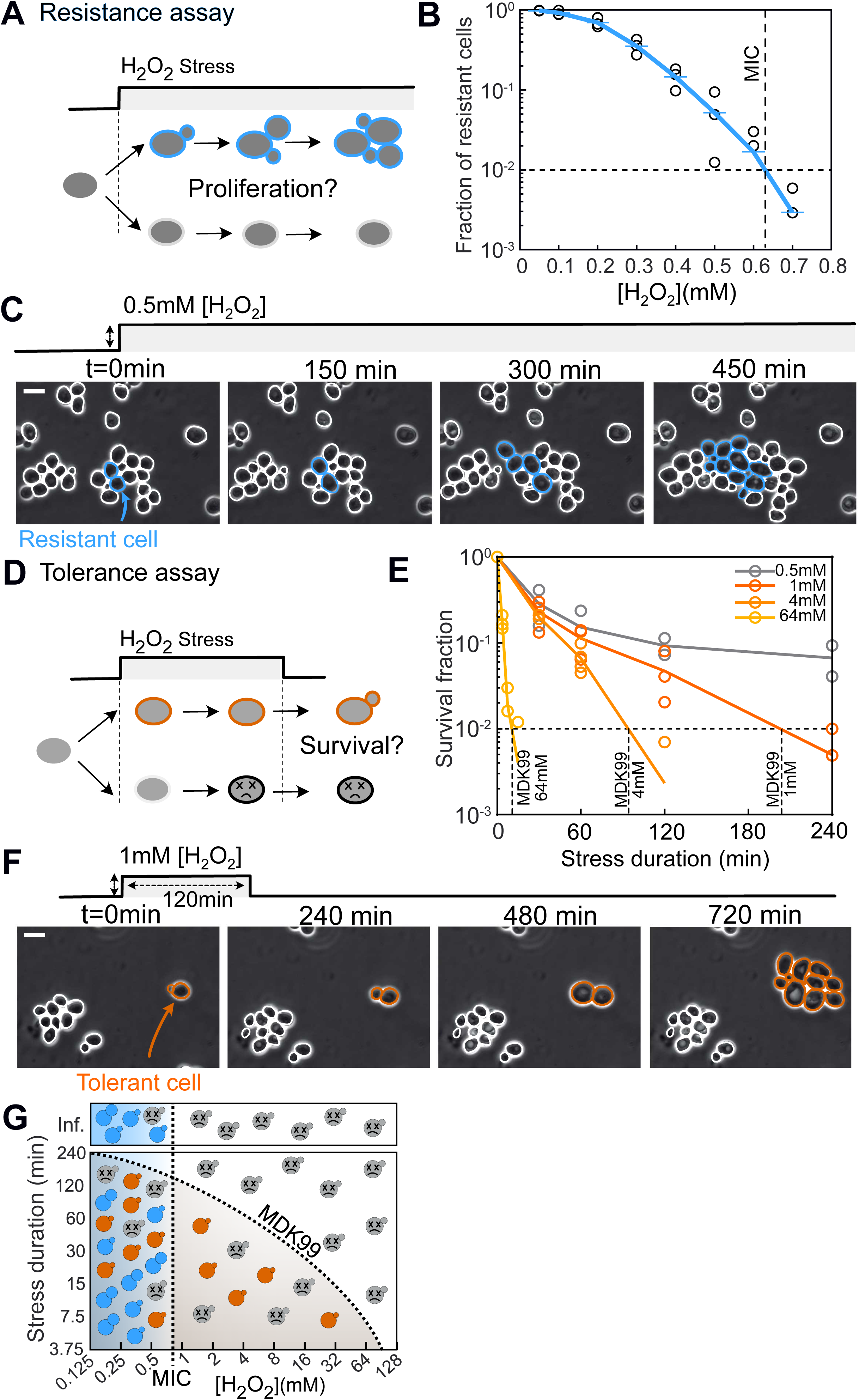
Resistance and tolerance are distinct properties of the response to hydrogen peroxide. **(A)** Diagram of the resistance assay. Blue (resp. white) cell contours indicate resistant (resp. non resistant) cells. **(B)** Fraction of resistant cells at different H_2_O_2_ concentrations. The vertical dashed line indicates the Minimal Inhibitory Concentration (MIC). Each open circle corresponds to a technical replicate. The blue curve indicates the mean fraction over 3 technical replicates per stress condition (n>100 cells for each replicate). **(C)** Representative time series of cells growing in the microfluidic device during a resistance assay at the indicated concentration. Blue (resp. white) cell contours indicate resistant (resp. non resistant) cells. **(D)** Diagram of the tolerance assay. Orange cell contours indicate tolerant cells (i.e. cells that resume growth after stress release), while dark gray contours represent dead cells. **(E)** Survival fraction of WT cells for different stress duration and at different concentrations. Each line represents the mean survival fraction based on technical replicates (represented as open circles, N = 2 or 3 per condition, n>100 for each replicate). Data points lower than 10^−3^ are not represented. **(F)** Representative time series of cells growing in the microfluidic device. Cells were exposed to a 1 mM H_2_O_2_ stress during 120min. Orange (resp. white) contours indicate tolerant (resp. non tolerant, or dead) cells. **(G)** Sketch showing the fate of cells (blue: resistant; orange: tolerant; gray: dead) as a function of H_2_O_2_ stress duration and concentration. The dashed line indicates the MIC and the MDK99.

In the second experiment, we measured cell survival (i.e., tolerance assay; see Figures 1D and 1F) by quantifying the fraction of cells present at the onset of stress exposure that recovered proliferation after a specific stress duration, excluding daughter cells born during stress to avoid accounting for resistance. Plotting the fraction of tolerant cells against the stress duration (Figure 1E and S1B) revealed a sharp decline in survival with increasing durations. This decay rate was [H_2_O_2_]-dependent and quantified using the MDK_99_ (Minimal Duration to Kill 99% of the population; Figure S1A).

Notably, this effect was independent of the cell cycle phase, as both budding and non-budding cells showed comparable survival fractions above the MIC (Figure S1C and S1D). Below the MIC (gray curve, 0.5 mM in Figure 1E), cell survival declined sharply for shorter stress durations but plateaued for longer durations, with survival fractions of 0.067 and 0.057 after 4- and 8-hour exposures, respectively. This plateau suggests a coexistence of distinct cell fates — resistance, tolerance, and mortality — at intermediate stressor concentrations.

Importantly, above the MIC, cells could survive brief stress exposure even though they were unable to proliferate at that concentration. For instance, ∼10% of the population survived a 1-hour exposure to 1 mM H_2_O_2_ (Figure 1E), despite fewer than 1% of cells being able to grow at this concentration (Figure 1B). Additionally, a small fraction of cells survived brief exposures to concentrations nearly two orders of magnitude higher than the MIC (survival fraction >0.01 after 7.5 minutes at 64 mM; Figure 1E).

Taken together, these experiments indicate that survival under stress does not necessarily imply the ability to proliferate. Resistance and tolerance are thus distinct, measurable properties of adaptation to hydrogen peroxide. However, substantial heterogeneity in cell fates is observed within populations exposed to stress, particularly at intermediate stress concentrations (i.e., 0.5 mM), as summarized in Figure 1G.

### The Yap1-mediated transcriptional stress response is required for H_2_O_2_ resistance but not for H_2_O_2_ tolerance

According to the homeostatic framework, cell adaptation to hydrogen peroxide is partially mediated by activation of the Yap1 regulon (Figure 2A). To investigate whether and how this homeostatic response is linked to resistance, tolerance, or both, we used the Yap1-dependent transcriptional reporter Srx1pr-GFP-degron as a proxy for Yap1 regulon activation. Specifically, we quantified its mean expression one hour after stress exposure in response to increasing H_2_O_2_ doses (SRX1 encodes sulfiredoxin, a Yap1-regulated gene; Figure S2A-C, Figure 2A-B). We found that its expression increased linearly with [H_2_O_2_] at low doses (<0.2 mM; see the top panel in Figure 2C) but progressively decreased to zero at concentrations above 0.3 mM, thereby validating the use of this reporter. To determine how this transcriptional activation relates to resistance and tolerance, we analyzed cell-to-cell heterogeneity in transcriptional response and cell fate within the population. We defined a threshold of Srx1pr-GFP-degron fluorescence to identify cells capable of mounting a transcriptional response (referred to as “responders” hereafter; see Figure 2B and Methods). As expected, the fraction of responders decreased with [H_2_O_2_] concentrations above 0.3 mM (bottom panel in Figure 2C).

**Figure 2:**
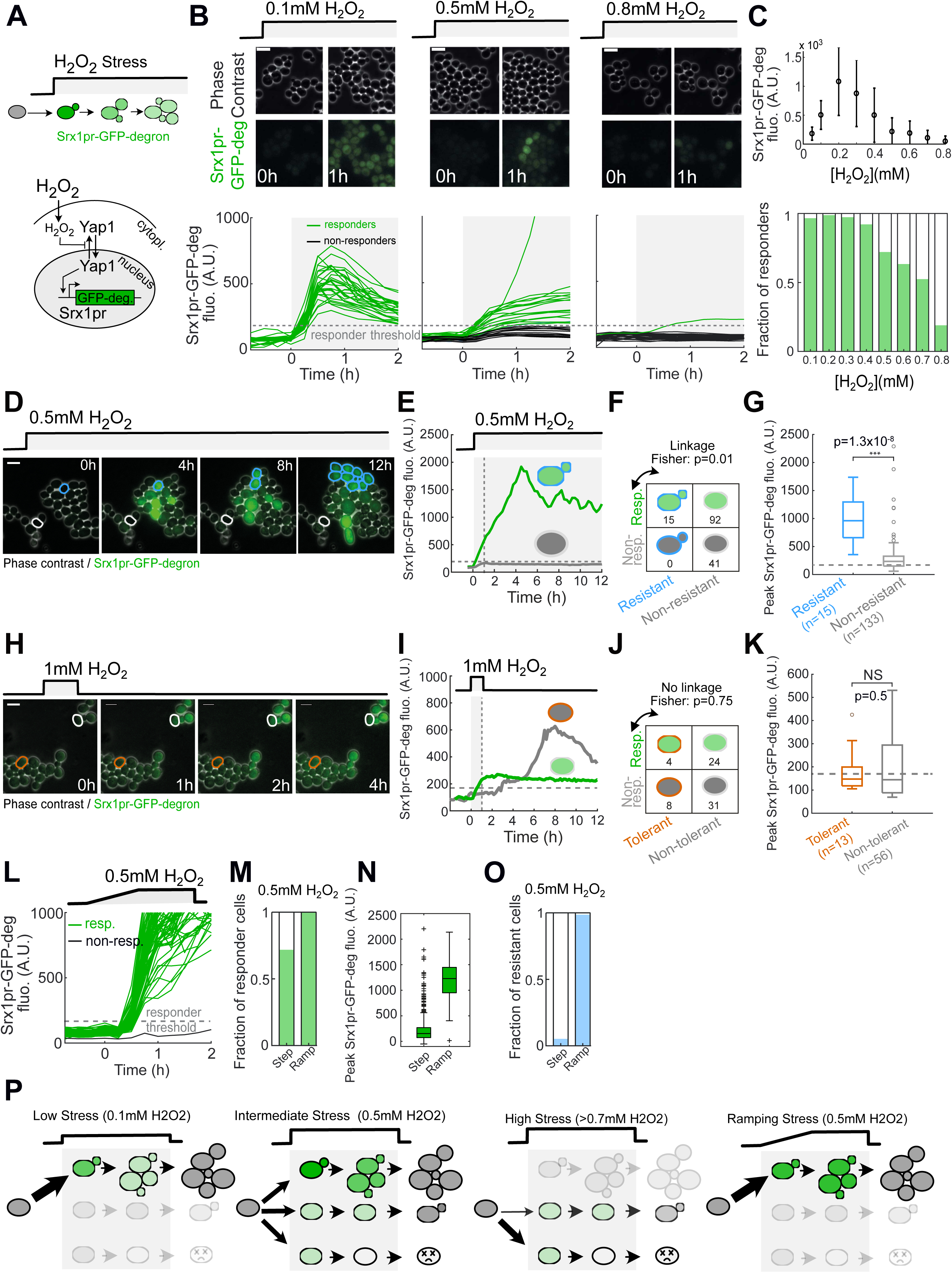
The Yap1-mediated transcriptional stress response is required for H_2_O_2_ resistance but not for H_2_O_2_ tolerance. **(A)** Sketch of the assay and representation of the Yap1-mediated transcriptional response for the Srx1 gene. **(B)** Top: Sequence of phase contrast and fluorescence images of a SRX1pr-GFP-deg reporter strain exposed to various constant H_2_O_2_ concentrations. Scale bar : 6.2 microns. Bottom: Sample single-cell GFP fluorescence quantifications as a function of time for responding and non-responding cells (green and black lines, respectively). The gray dashed line represents the responder threshold; N>100 cells for each condition. **(C)** Top: Mean expression of Srx1pr-GFP-degron (calculated at t=1h) as a function of H_2_O_2_ concentration. Error bars represent the standard error of the mean (n>100 cells for each condition). Bottom: fraction of responding cells as a function of H_2_O_2_ concentration (n>500 single-cells, pooled from at least N=2 technical replicates). **(D)** Sequence of overlaid phase contrast and fluorescence images of WT cells carrying an Srx1pr-GFP-deg reporter. Blue (resp. white) cell contours indicate typical resistant (resp. non-resistant) cells. **(E)** Sample single-cell trajectories of Srx1-GFP-deg cells for responding cells (green line) and non-responding cells (grey line). The gray dashed line indicates the “responder” threshold (see Methods for detail). The vertical dashed line (t=1h) indicates the time when the responding status of the cells is assessed. **(F)** Contingency matrix showing the number of responding and non-responding cells for both resistant and non-resistant cells (0.5 mM H_2_O_2_). The fisher test shows that the two binary variables are linked (p=0.01). **(G)** Boxplot of the Srx1pr-GFP-deg peak response (at t=1h following stress addition) in resistant (n=15) versus non-resistant cells (n=133). Statistical differences (two-sided Wilcoxon rank sum test) are indicated by *** for *p* < 0.001, NS for *p* > 0.05. **(H and I)** Same as (D and E), but for the tolerance assay (1mM H_2_O_2_, 60min). The orange contours indicate tolerance cells. **(J)** Contingency matrix showing the number of responding and non-responding cells for both tolerant and non-tolerant cells (1 mM H_2_O_2_). The fisher test shows that the two binary variables are not linked (p = 0.75). **(K)** Boxplot of the Srx1pr-GFP-deg peak response (at t=1h following stress addition) in resistant (n=13) versus non-resistant cells (n=56). Statistical differences (two-sided Wilcoxon rank sum test) are indicated by *** for *p* < 0.001, NS for *p* > 0.05. **(L)** Quantification of mean single-cell GFP fluorescence as a function of time for responding and non-responding cells (green and black lines, respectively) during a ramping stress. **(M)** Fraction of responder cells in response to step and ramp assays (0.5 mM H_2_O_2_); n=142 for ramp, step data are taken from Figure 2D-G for comparison. **(N)** Mean Srx1pr-GFP-deg expression one hour after a 0.5 mM H_2_O_2_ ramping exposure (step assay data are taken from Figure 2D-G for comparison). **(O)** Fraction of resistant cells in response to step and ramp assays (0.5 mM H_2_O_2_). n=142 for ramp, step data are taken from Figure 2D-G for comparison. **(P)** Sketch summarizing the fates of cells submitted to varying concentrations and temporal patterns of H_2_O_2_., along with their links to the transcriptional response. Filled green cells represent responding cells. Light-gray cells are dead. The black arrows indicate the respective fractions of the cells undergoing a particular fate (top: resistant; middle: tolerant; bottom: dead).

Importantly, we found that all resistant cells were responders (Figures 2D-F), indicating that the transcriptional activation of the Yap1 regulon is necessary for resistance. Furthermore, a contingency analysis revealed a significant association between transcriptional response and the resistance phenotype (Fisher’s exact test, p = 0.01; see Figure 2F). This association was further confirmed by showing that resistant cells displayed significantly higher antioxidant expression compared to non-resistant cells (Figure 2G). In contrast, no such correlation was observed between transcriptional activation and tolerance following exposure to 1 mM H_2_O_2_ (Figures 2H-K). Taken together, these results suggest that the Yap1-mediated transcriptional response is required for resistance but not for tolerance.

If the transcriptional response is crucial for resistance, any intervention that facilitates its activation should benefit cell proliferation under stress. Therefore, a gradual increase in stress (ramping stress) should allow for greater resistance than an abrupt stress exposure, as it provides more time for the homeostatic system to respond. During a ramp with an amplitude of 0.5 mM, the vast majority of cells displayed Srx1pr-GFP-degron expression above the “responder” threshold (Figures 2L-N) and maintained resistance (Figure 2O), thus confirming our hypothesis. This observation stood in stark contrast to the phenotypes observed after an abrupt increase to 0.5 mM H_2_O_2_ (step stress, see Figures 2M-O).

Overall, these experiments contribute to linking the activation of the Yap1 regulon to the different cell fates observed upon exposure to H_2_O_2_, as summarized in Figure 2P. Under low stress conditions (∼0.1 mM H_2_O_2_), Yap1-mediated gene expression supports cell proliferation. At intermediate concentrations (∼0.5 mM), non-resistant cells emerge in the population, exhibiting either tolerance to stress or cell death, regardless of their ability to activate the transcriptional response. These non-resistant cells become dominant in the population beyond the MIC (>0.7 mM). Finally, ramping stress improves the ability of cells to activate their transcriptional response and resist. Hence, from a practical perspective, ramping stress provides an effective experimental approach to assess the capacity of cells to resist and/or tolerate stress while maintaining a quasi-steady state. In contrast, the acute stress protocol evaluates components of the response that are independent of transcription.

### Investigating the genetic determinism of resistance and tolerance unravels a trade-off between stress resistance and stress tolerance

We then sought to investigate the genetic determinants of resistance and tolerance. Based on the above results, we conducted a genetic screen using mutant strains deleted for one or more genes previously described as important for the H_2_O_2_ stress response. This included genes regulated by Yap1, as well as mutants associated with the general stress response (see Figure S3A for the list of mutants). To assess cell resistance, we applied a stress ramping protocol and measured the relative fold-change in biomass production of each mutant compared to the wild type (WT) following a 4-hour exposure to 0.5 mM H_2_O_2_ (see Figures 3A, 3B, S3B, and S3C and Methods). As shown in Figure 2, this protocol evaluates the capacity of cells to sustain growth independently of the transient acute response observed during stepwise stress exposure. Conversely, we used a pulsed stress protocol (4 hours at 0.5 mM) to measure post-stress cell survival as a proxy for tolerance (Figures 3C, 3D, S3B, and S3D).

**Figure 3:**
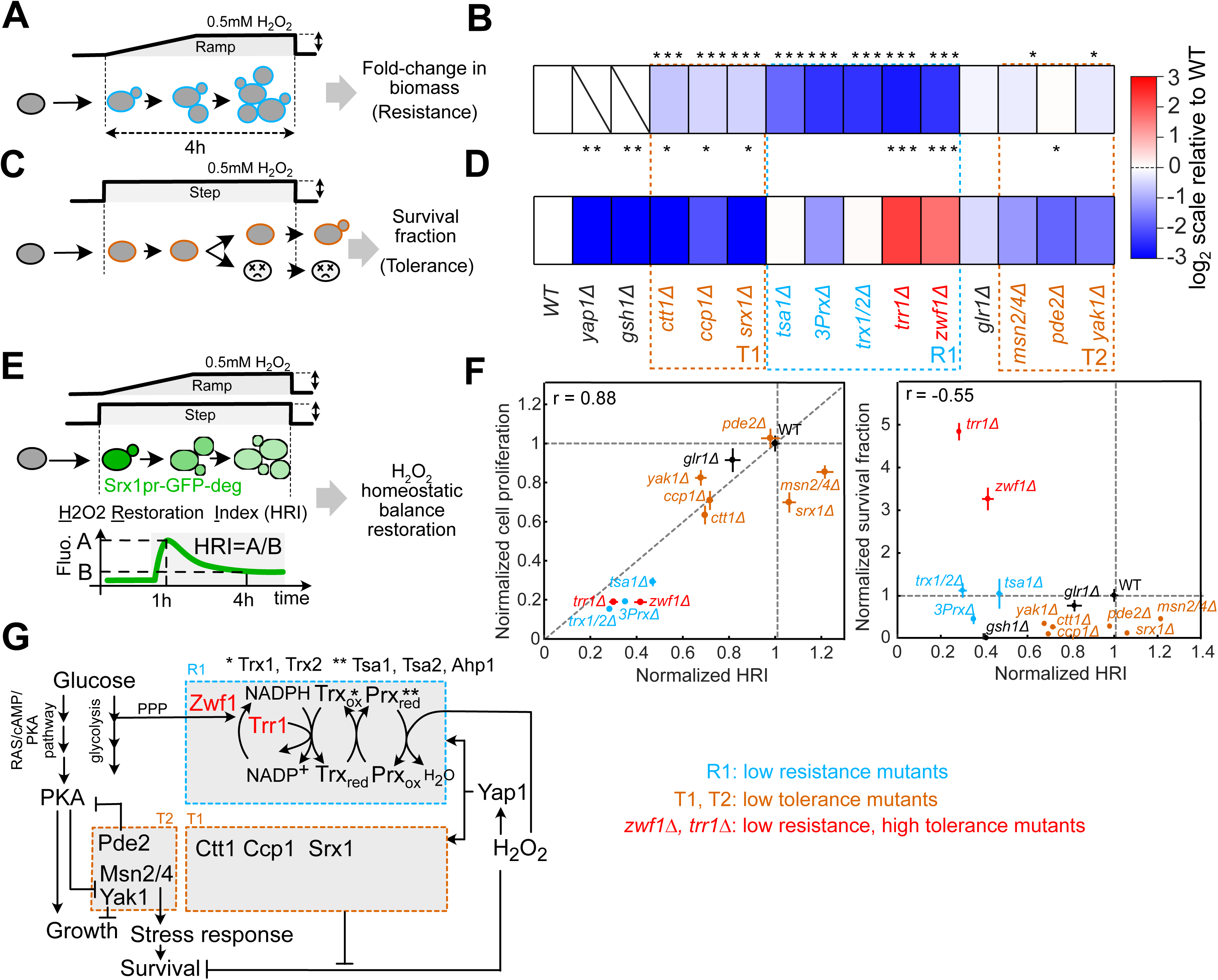
Investigating the genetic determinism of resistance and tolerance unravels a trade-off between the two defense strategies. **(A)** Sketch of the assay performed to measure cellular resistance to H_2_O_2_ in the collection of mutants. Resistance is assessed by measuring the fold-change in total biomass production between t=0h and t=4h during a ramping stress assay for cells initially present at t=0h. **(B)** Quantification of the normalized cell proliferation in the ramp assays for the indicated mutants (between n = 6 and 55 micro-colonies depending on mutants, from N = 3 different technical replicates, see Methods and Source data file for details). Stars indicate the result of a non-parametric Wilcoxon-Mann-Whitney statistical test comparing the indicated mutants to the WT. **** for *p* < 0.0001, *** for *p* < 0.001, ** for *p* < 0.01, * for *p* < 0.05, NS for *p* > 0.05. **(C-D)** Same as (A-B), but for the tolerance assay. N = 3 technical replicates for each condition. The dashed blue rectangle R1 represents a group of mutants with low resistance. The dashed orange rectangles T1 and T2 represent two groups of mutants with low tolerance. Stars indicate the result of a t-test comparing the indicated mutants to the WT. **** for *p* < 0.0001, *** for *p* < 0.001, ** for *p* < 0.01, * for *p* < 0.05, NS for *p* > 0.05. **(E)** Sketch explaining the quantification of the H_2_O_2_ restoration index (HRI), defined as the ratio between the Srx1pr-GFP-deg fluorescence at t=1h to that measured at t=5h during indicated stress patterns. **(F)** Left: normalized cell proliferation during ramps (from (**B**)) as a function of the H_2_O_2_ restoration index (HRI), normalized to WT; r indicates the Pearson correlation coefficient. HRI are measured with n>50 single-cells for each condition. Error bars represent the s.e.m, between n=6 and 54 micro-colonies depending on mutants for cell proliferation and n>50 single-cells for HRI measurements. Right: same as the left panel, but for the survival fraction (data taken from (D)). Error bars represent the s.e.m, N=3 technical replicates for survival fractions and n>50 single-cells for HRI measurements. **(G)** Sketch showing the mechanistic links between mutants assayed in Figure 3. The R1 box indicates the low resistance mutants identified in (B); The T1 and T2 boxes show the two groups of low tolerance mutants, related to panel (D).

To validate this methodology, we first checked that *yap1*Δ and the *gsh1*Δ mutants (*GSH1* encodes an enzyme involved in the biosynthesis of glutathione) exhibited no proliferation or tolerance under both stepping and ramping stress protocols. This observation is consistent with their essential roles in maintaining general redox homeostasis and other canonical cellular functions^23^, see Figure 3B, 3D, and S3E. Next, our analysis identified a group of mutants with a pronounced resistance defect compared to the wild type (WT), highlighted by the R1 rectangle in Figure 3B (see also Figure S3E).

This group included genes essential for maintaining a functional peroxidatic cycle (referred to as the Prx/Trx pathway; see schematic in Figure 3G), which is orchestrated by enzymes known as 2-Cys peroxiredoxins (i.e., Tsa1, Tsa2, and Ahp1; the triple mutant is referred to as 3PrxΔ in Figure 3B). Peroxiredoxins catalyze the reduction of H_2_O_2_ and are recycled by thioredoxins (Trx1 and Trx2), which are, in turn, reduced by the NADPH-dependent thioredoxin reductase (Trr1)^42^. NADPH is primarily synthesized through the pentose phosphate pathway, the first step of which is catalyzed by glucose-6-phosphate dehydrogenase (G6PDH), encoded by the ZWF1 gene^43^. Importantly, all these mutants — and combinations thereof — exhibited a consistent decrease in resistance (Figure 3B).

In contrast, deletion of the cytoplasmic catalase (*CTT1*), cytochrome c peroxidase (*CCP1*), or sulfiredoxin (*SRX1*), which reduces the hyperoxidized form of peroxiredoxins, only slightly affected cell resistance compared to mutants of the Prx pathway (Figure 3B). Similarly, inhibiting the general stress response by deleting the *MSN2/4* transcription factors, or the yeast kinase *YAK1*, which partially mediates PKA-dependent activation of Msn2/4^44^, or hyperactivating the Protein Kinase A (PKA) pathway by deleting *PDE2* — both interventions previously shown to increase sensitivity to H_2_O_2_^37,45^ — did not result in a significant reduction in resistance.

Instead, all the corresponding knockout strains exhibited a significant decrease in cell tolerance compared to the WT under stepping stress conditions (see T1 and T2 groups in Figures 3B and 3D, see also Figure S3E). Hence, this analysis demonstrates that, among the genes involved in the oxidative stress response, some are essential for restoring or maintaining proliferation, while others primarily contribute to cell survival.

We then hypothesized that cells with decreased resistance would be impaired in their ability to restore a physiological internal H_2_O_2_ balance upon stress exposure. To test this, we used the Srx1pr-GFP-degron reporter as a proxy for internal H_2_O_2_ balance, assuming that the decline in fluorescence following peak expression accurately reflects the deactivation of Yap1 transcription due to a drop in internal H_2_O_2_ concentration, as previously described^7^. We integrated this reporter into all the mutants mentioned above and quantified the “H_2_O_2_ Restoration Index” (HRI) for each strain in response to a 0.1 mM H_2_O_2_ step stress (see Figure 3E and Methods). The HRI was defined as the ratio between the mean cytoplasmic fluorescence at t = 1 hour (corresponding to the peak Srx1pr-GFP-degron level in the WT under these conditions) and t = 5 hours (see Figure 3E and S4A-C). To further validate our methodological approach, we quantified the HRI using the cytoplasmic H_2_O_2_ sensor Hyper7^46^ in the WT (Figure S5A and S5B) and eight mutants from the screen (Figure S5C). We found a strong correlation between the HRI calculated using either the Srx1pr-GFP-degron reporter or the Hyper7 sensor (r = 0.76, Figure S5D), reinforcing our hypothesis that the HRI provides a reliable measure of H_2_O_2_ balance recovery across different mutants, with higher HRIs indicating faster recovery following H_2_O_2_ exposure. By normalizing the HRI of each mutant to that of the WT, we observed that poorly resistant mutants (R1 group) exhibited a reduced HRI (left panel in Figure 3F). Overall, we found a strong positive correlation between HRI and resistance (r = 0.88, left panel in Figure 3F), confirming the link between the ability to degrade H_2_O_2_ and the capacity to maintain cell proliferation under stress. In contrast, the HRI in tolerant mutants from the T1 and T2 groups showed a weaker, negative correlation with tolerance (r = −0.55, right panel in Figure 3F).

Apart from the previously mentioned gene clusters, our attention was drawn to two specific mutants: *zwf1Δ and trr1Δ*. These mutants exhibited diminished resistance (Figure 3B) and a low HRI (Figures 3F, S4C, and S5D), aligning with other factors in the R1 cluster. However, we observed a remarkable 3- and 5-fold increase in tolerance compared to the WT, respectively (Figures 3D). This finding was unexpected, as these mutants had previously been described as sensitive to H_2_O_2_ in various model organisms^12,33,47–50^. Therefore, our analysis of candidate genes allowed us to categorize mutants into distinct functional groups, thereby refining the role of the corresponding genes in orchestrating adaptation to H_2_O_2_. Specifically, we identified two mutants, *zwf1Δ* and *trr1Δ*, in which a pronounced defect in H_2_O_2_ resistance, along with impaired H_2_O_2_ homeostasis, was unexpectedly associated with a hyper-tolerant phenotype (Figure 3G).

### Proliferation and survival are antagonistic features of the oxidative stress response

We then sought to characterize further how the mutation of the *ZWF1* gene induces both a decline in proliferation under H_2_O_2_ stress and an increase in tolerance, and whether this effect depends on the internal H_2_O_2_ balance.

First, complementary stress response assays revealed that the MIC of the *zwf1*Δ mutant was significantly lower than that of the WT (0.3 mM and 0.7 mM, respectively; Figure 4A). To determine whether this drop in resistance was due to an impaired transcriptional response of the Yap1 regulon, we quantified the expression of the Srx1pr-GFP-degron reporter in response to mild H_2_O_2_ concentrations ranging from 0.05 to 0.2 mM (i.e., below the MIC of the *zwf1Δ* mutant). Our results indicated that the mutant displayed a maximal expression level similar to or higher than that of the WT (see left and middle panels in Figure 4B, and Figure S7A), suggesting that its resistance defect is not due to a lack of regulon activation. Instead, unlike the WT, these experiments confirmed that the mutant fails to restore H_2_O_2_ balance, as evidenced by the persistent activation of the Srx1pr-GFP-degron reporter (see left and middle panels in Figure 4B) and the nuclear localization of the Yap1-sfGFP fusion protein (Figures S6B and S6C).

**Figure 4:**
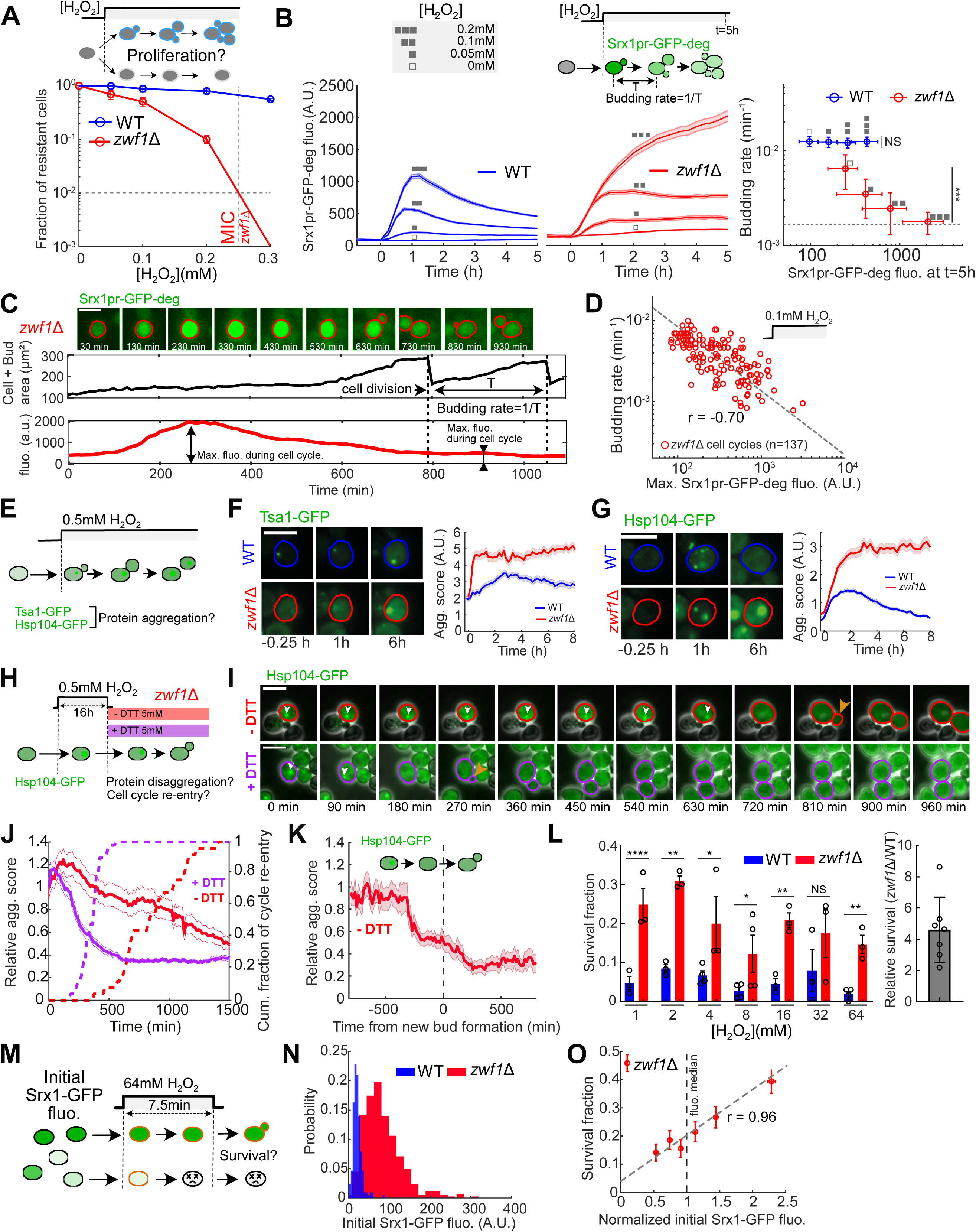
Proliferation and survival are antagonistic features of the oxidative stress response. **(A)** Top: sketch of the stress response assay used to compare WT and *zwf1*Δ resistance; Bottom: fraction of resistant cells in WT (blue) and *zwf1*Δ (red) cells as a function of H_2_O_2_ concentration (N = 3 independent replicates, n>100 for each replicate). The vertical and horizontal dashed lines indicate the MIC for the *zwf1*Δ mutant. **(B)** Top: H_2_O_2_ concentrations used in the assay and sketch of the variables (Srx1pr-GFP-degron level and budding rate) measured in the assay; Bottom left and middle: quantification of the mean Srx1pr-GFP-deg fluorescence expression +/− s.e.m. over time in WT (blue lines) and *zwf1*Δ mutant (red lines) in response to the indicated H_2_O_2_ concentrations (n>100 at stress beginning for each concentration). Bottom right: mean budding rate of resistant cells +/− one standard deviation (n>33 single-cells per condition) as a function of the mean Srx1pr-GFP-deg fluorescence expression after 5h under stress +/− one standard deviation in WT (blue, n>100) and zwf1Δ mutant (red, n>100). Statistical differences in budding-rates from 0 to 0.2 mM were evaluated for WT and *zwf1*Δ strains using a one-way ANOVA test for multiple comparisons. **** for *p* < 0.0001, *** for *p* < 0.001, ** for *p* < 0.01, * for *p* < 0.05, NS for *p* > 0.05. **(C)** Top panel: sequence of overlaid phase contrast and Srx1pr-GFP-deg fluorescence of a representative *zwf1Δ* cell during one cell cycle. The red line delineates the cell contour Middle/bottom panels: quantification of cell and bud area (black line, top panel) as well as Srx1pr-GFP-degron fluorescence (red line, bottom panel) over time. The black dashed lines indicate the time of bud emergence. **(D)** Budding rate of individual *zwf1*Δ cells exposed to 0.1 mM for more than 5h as a function of the maximal fluorescence expression of the SRX1pr-GFP-deg reporter during the corresponding cell cycle (n=137). *r* is the correlation coefficient between the two quantities. **(E)** Sketch of the assay to measure protein aggregation during H_2_O_2_ step stress exposure. **(F)** Left: sequence of Tsa1-GFP fluorescence images for WT and *zwf1Δ* cells at indicated times; Colored lines delineate cell contours; Right: quantification of the aggregation score +/− s.e.m. of Tsa1-GFP foci for the indicated strains (see Methods for details). **(G)** Same as **(F)**, but for the Hsp104-GFP reporter. **(H)** Sketch of the protein disaggregation assay in a *zwf1Δ* strain with a Hsp104-GFP marker, in the absence (red) or presence of DTT (magenta). **(I)** Time series showing phase contrast and GFP fluorescence images of *zwf1*Δ cells carrying an Hsp104-GFP reporter according to the protocol described in (H). Red and magenta contours indicate a cell of interest. White arrows indicate the Hsp104-GFP aggregates. The orange arrow marks the first bud observed after stress release. Scale bar 6.2 µm. **(J)** Left axis: Hsp104-GFP aggregation score +/− s.e.m., normalized to the aggregation score at t=0min (Solid red and magenta lines), following stress release according to the protocol in **(H)**. Only surviving cells are included in the analysis (see Methods, n>50 for each condition at all times). Right axis: cumulative fraction of cells re-entering the cell-cycle with or without DTT addition (red dashed line, n=35, and purple dashed line, n=50, respectively, see Methods for detail). **(K)** Hsp104-GFP aggregation score +/− s.e.m. after stress release in the population of surviving *zwf1Δ* cells, after synchronization from new bud formation for each single cell. **(L)** Left panel: mean survival fraction of WT (blue bars) and *zwf1*Δ (red bars) cells as a function of H_2_O_2_ concentration. For each concentration, the duration of stress exposure is set to half of the WT MDK99 (see also Figure S1A); N = 3 to 4 technical replicates (black open circles), n>100 for each replicate. Statistical differences (one-sided two-sample t-test) are indicated by **** for *p* < 0.0001, *** for *p* < 0.001, ** for *p* < 0.01, * for *p* < 0.05, NS for *p* > 0.05. Right panel: fold-change in *zwf1*Δ survival versus WT after pooling all tested concentrations in the left panel (N=7). **(M)** Sketch showing the protocol used to measure cell survival to a step stress as a function of the initial Srx1-GFP fluorescence level. **(N)** Distribution of basal fluorescence of the Srx1-GFP fusion protein in WT and *zwf1*Δ strains (n=807 cells) at the onset of a 7.5 min stress exposure at 64 mM H_2_O_2_. **(O)** Mean survival in *zwf1*Δ cells from panel (N) sorted by their normalized pre-stress Srx1-GFP signal +/− s.e.m (n=135 cells for group 1 to 5 and n= 132 cells for group 6). Data were pooled into 6 bins according to the initial Srx1-GFP level of the cells. *r* is the correlation coefficient between both quantities. The vertical dashed line indicates the median fluorescence in the population. The other dashed line shows linear fit to the data.

Notably, the reduction in mean growth rate at steady state was negatively correlated with the prolonged activation of the Yap1 regulon (assessed at t = 5 hours after the onset of stress exposure; see the right panel in Figure 4B and see also Figure S6D, and S6E). This negative correlation was also observed at the single-cell level, where individual dividing cells experienced erratic arrests in their cell cycle, concomitant with bursts of Srx1pr-GFP-degron expression (Figures 4C and 4D). Altogether, these results suggest that the *zwf1Δ* mutation causes a pronounced, yet heterogeneous and fluctuating, H_2_O_2_ imbalance that, in turn, leads to reduced cell proliferation.

To further investigate the impact of H_2_O_2_ imbalance on cellular function, we assessed the level of proteome oxidation in the *zwf1Δ* mutant. First, we monitored the formation of fluorescent foci of the Tsa1-GFP fusion protein in cells exposed to 0.5 mM H_2_O_2_ over 8 hours (Figure 4E). Tsa1 is known to form supramolecular assemblies when hyperoxidized in response to severe H_2_O_2_ stress, serving as a marker of oxidative damage^51,52^. In the WT, we observed a progressive appearance of Tsa1-GFP foci in response to stress, followed by a decrease likely due to stress adaptation (Figure 4F). In contrast, in the *zwf1Δ* mutant, aggregation occurred much faster and was irreversible over the 8-hour stress exposure. We obtained similar results using the Hsp104-GFP fusion protein (Figure 4G), which serves as a general marker of protein aggregation. Hsp104 is a disaggregase that binds misfolded, aggregated proteins and forms localized foci in response to H_2_O_2_ stress^53^. These experiments provided evidence that the H_2_O_2_ imbalance in the *zwf1*Δ mutant drives the stable appearance of hallmarks of proteome oxidation.

Next, to determine how protein oxidation is linked to cell proliferation, we released cells previously exposed to 0.5 mM H_2_O_2_ for 16 hours into a stress-free medium and monitored the dynamics of protein aggregation using the Hsp104-GFP reporter (Figure 4H). Strikingly, we observed a progressive disaggregation of Hsp104-GFP foci (Figures 4I and 4J) that temporally coincided with cell cycle re-entry in cells that survived the stress (Figures 4K). This observation suggested that recovery of cell proliferation requires proteome reduction. Supporting this hypothesis, the addition of 5 mM DTT (a reducing agent) to the medium upon H_2_O_2_ stress release significantly accelerated both the disappearance of protein aggregates and cell cycle re-entry (Figure 4I-J). Altogether, these experiments demonstrate that protein oxidation observed during stress exposure in the *zwf1*Δ mutant is closely associated with proliferation arrest. However, this oxidation is reversible and does not impair the cells’ ability to resume growth after stress removal. Hence, these findings suggest that, like internal H_2_O_2_ imbalance, proteome oxidation is not necessarily detrimental to cell survival in the *zwf1Δ* background.

To further characterize the enhanced ability of the *zwf1*Δ mutant to survive H_2_O_2_ stress exposure, we compared its survival to that of the WT in cells subjected to increasing doses of H_2_O_2_ above the MIC (ranging from 1 mM to 64 mM) for durations equal to half of the MDK_99_ at each indicated concentration (Figure 4L, also see Figure S1B). We confirmed a mean fold-change in survival of 4.6 ± 2.1 across all conditions compared to the WT strain (right panel in Figure 4L), confirming the mutant’s superior tolerance. We verified that this gain of tolerance was not due to an increased proportion of non-budding or quiescent cells in the mutant (Figure S1C and S1D) and was not restricted to a specific strain background (Figure S1E and S1F), suggesting the involvement of a more specific mechanism. Finally, we leveraged the large intrinsic cell-to-cell variability in H_2_O_2_ balance within the *zwf1*Δ mutant to investigate how the initial H_2_O_2_ imbalance affects cell survival after a severe stress pulse (Figure 4M). To do this, cells were exposed to 64 mM H_2_O_2_ for 7.5 minutes, and the fraction of surviving cells was scored as a function of their initial Srx1-GFP expression level (Figure 4N). Strikingly, we found that the initial imbalance in internal H_2_O_2_ was a strong predictor of survival to future acute stress exposure (r = 0.96 for binned data; see Figure 4O), with cells displaying a large positive imbalance showing improved stress tolerance.

Altogether, this analysis reveals a trade-off between H_2_O_2_ resistance and tolerance in the *zwf1Δ* mutant compared to the WT. It suggests the existence of a mechanism that signals H_2_O_2_ imbalance to shut down growth control and trigger protective effects on cellular function, despite high internal oxidation levels.

### Stress tolerance requires the Prx/Trx pathway to drive PKA inhibition

Genetic perturbations that sustain high PKA activity during stress exposure have long been associated with increased stress sensitivity^35^. In line with this, our candidate-gene approach confirmed that both the general stress response and PKA inhibition are specifically required for H_2_O_2_ tolerance (see *msn2/4*Δ and *pde2*Δ mutants in Figure 2B). Given this, we next asked whether PKA regulation contributes to the trade-off between low resistance and high tolerance observed in the *zwf1*Δ mutant.

First, we hypothesized that *ZWF1* deletion could lead to down-regulation of PKA activity, thereby reducing cell proliferation and enhancing tolerance. To test this, we monitored the nuclear shuttling of an Msn2-GFP fusion protein as a readout of PKA activity (Figure 5A), as previously described^37^. Msn2-GFP remained nuclear at all times in the *zwf1*Δ mutant during stress exposure, consistent with strong PKA inhibition (Figures 5B-D). In contrast, the transient nuclear relocation of Msn2-GFP was greatly reduced in the *pde2*Δ strain compared to the WT, consistent with the previously described constitutive PKA activation in this background (Figures 5B-D). Importantly, introducing the *pde2*Δ mutation into the *zwf1*Δ background caused Msn2-GFP to remain fully cytoplasmic during stress exposure (Figures 5B-D), suggesting that PKA was indeed reactivated in this strain upon H_2_O_2_ exposure. This effect was further amplified by supplementing the medium with cAMP (Figure 5B-D), a procedure known to exacerbate the pde2Δ phenotype^54^. Therefore, *ZWF1* deletion inhibits PKA, but this effect can be overridden by forced PKA activation.

**Figure 5:**
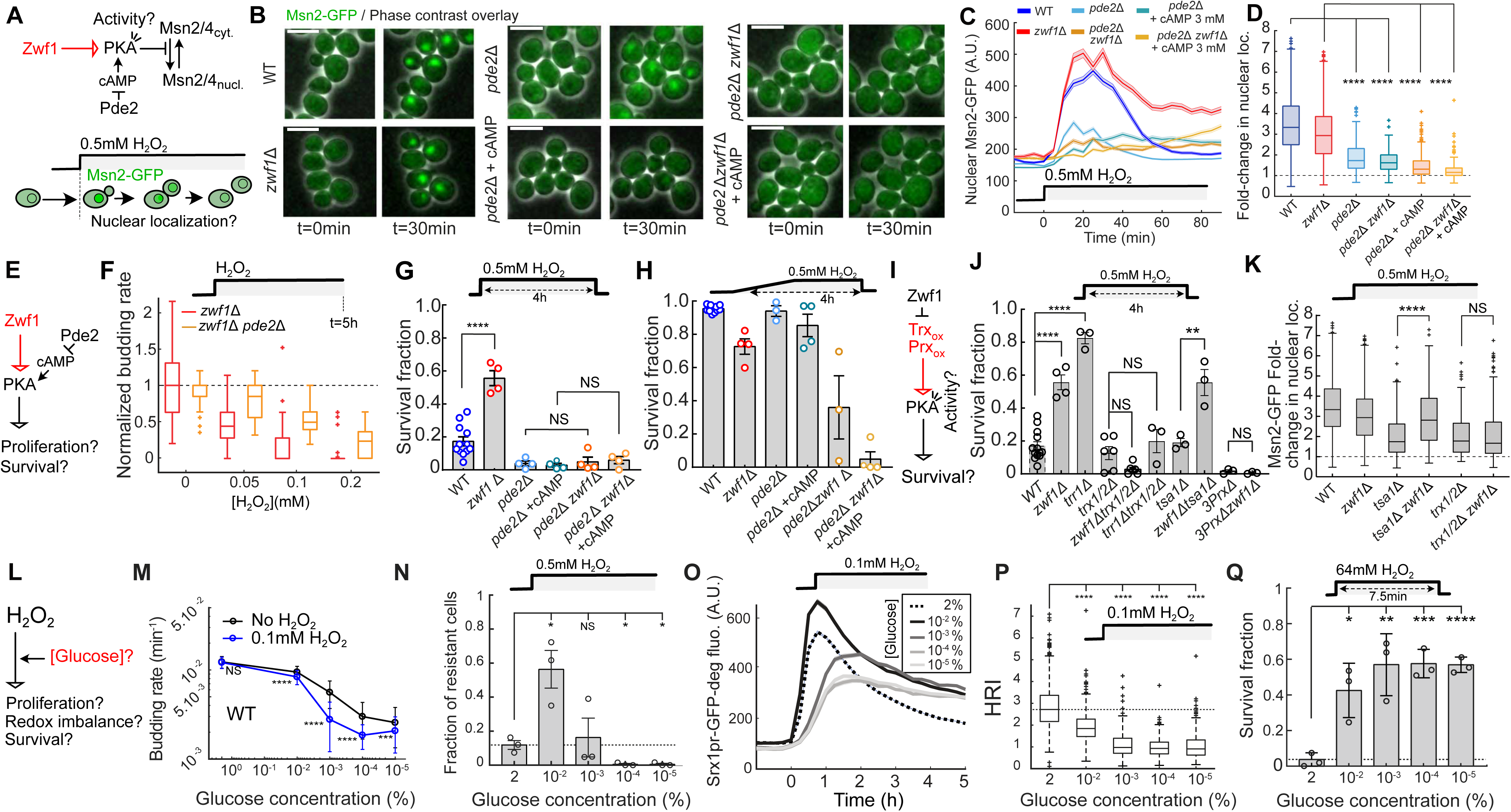
Stress tolerance requires thioredoxins to drive PKA inhibition. **(A)** Schematic illustrating the mechanistic link between the Pde2, PKA and Msn2 and raising the hypothesis of a genetic interaction of this pathway with *ZWF1*. **(B)** Overlays of phase-contrast and fluorescence images of cells carrying an Msn2-GFP fusion before (t=0min) and after (t=30min) the addition of 0.5 mM H_2_O_2_ for the indicated strains. Scale bar: 6.2µm. **(C)** Nuclear localization score of Msn2-GFP over time +/− s.e.m. for the identical strain backgrounds. Sample size: n = 162 (WT), 242 (*zwf1*Δ), 139 (*pde2*Δ), 179 (*pde2*Δ + cAMP), 184 (*zwf1*Δ*pde2*Δ) and 184 (*zwf1*Δ*pde2*Δ + cAMP). **(D)** Boxplots of the fold-change in the Msn2-GFP nuclear score between the control condition and 30 min after stress the addition of a 0.5mM H_2_O_2_ concentration, for each indicated strain background (n>100 for each condition). Statistical analysis based on a two-sided Mann-Whitney U test. **(E)** Schematic illustrating the potential interaction between *ZWF1* and *PDE2* during H_2_O_2_ stress exposure, as investigated in panels (F-H). **(F)** Boxplot of the normalized budding rate of cells exposed to the indicated H_2_O_2_ concentration. For each strain, budding rates were normalized between the median budding rate without stress (0 mM condition) and the minimal budding rate experimentally considered, 0.0017 min^−1^. Sample size between 31 and 105 single-cells for budding rate quantifications. **(G-H)** Survival fraction (gray bars) in response to a 4 h 0.5 mM H_2_O_2_ step (**G**) or ramp (**H**) exposure for the indicated strain backgrounds. Bars represent means of independent technical replicates (open circles, N ≥ 3 technical replicates per condition, n>100 for each replicate). Error bars indicate the s.e.m. The statistical analysis in G is based on a one-sided t-test. **(I)** Schematic illustrating the potential interaction between *ZWF1*, Thioredoxins (Trx), Peroxireodxins (Prx) and *PDE2* during H_2_O_2_ stress exposure, as investigated in panels (J) and (K). **(J)** Same as (G), but shown for different strain backgrounds. **(K)** Same as (D), but shown for different strain backgrounds. **(L)** Schematic illustrating the potential role of glucose concentration on resistance, redox imbalance, and tolerance. **(M)** Mean budding rate of WT resistant cells +/− std without (black line, n>62 cells per data point) or with 0.1 mM H_2_O_2_ (blue line, n>61 per glucose concentration), measure 5 h after stress induction, as a function of glucose concentration. The statistical analysis is based on a two-sided Man Whitney U test. **(N)** Fraction of resistant cells exposed to a 0.5 mM H_2_O_2_ stress. Open circles are independent technical replicates (N = 3 with n>100 for each replicate); bars represent the mean value and the error bars are the s.e.m. The statistical analysis is based on a two-sided t-test. **(O)** Mean Srx1pr-GFP-deg cell fluorescence +/− s.e.m. over time in response to a 0.1 mM H_2_O_2_ stress (at t=0h) for different glucose concentrations. n>100 single-cells for each concentration. **(P)** H_2_O_2_ restoration index (HRI) under 0.1 mM H_2_O_2_ as a function of glucose concentration. The statistical analysis is based on a two-sided Man Whitney U test. n>100 single-cells for each concentration. **(Q)** Fraction of tolerant cells after a 64 mM H_2_O_2_ stress exposure for 7.5 min. Open circles are independent technical replicates (N = 3 with n>100 for each replicate), bars represent the mean value and the error bars are the s.e.m. The statistical analysis is based on a two-sided t-test. **(A-Q)** For all statistical analysis, significance levels are: **** for *p* < 0.0001, *** for *p* < 0.001, ** for *p* < 0.01, * for *p* < 0.05, NS for *p* > 0.05.

We then reasoned that constitutive PKA activation could rescue the impaired cell growth of the zwf1Δ mutant during stress exposure (Figure 5E). To test this hypothesis, we measured the budding rate of the double mutant at steady state, i.e., measured at t = 5 hours post-stress onset. Consistent with our hypothesis, deleting *PDE2* in the *zwf1Δ* background improved the overall proliferation rate during stress exposure, compared to the *zwf1Δ* single mutant (Figure S7A). Furthermore, the relative decline in proliferation was shifted towards higher H_2_O_2_ concentrations, demonstrating a net gain in resistance in the double mutant (Figure 5F). Importantly, the double mutant exhibited a steady-state activation level of the Srx1pr-GFP-degron reporter similar to that of the *zwf1Δ* mutant (Figure S7B), indicating that the gain in resistance due to PDE2 deletion is not attributable to H_2_O_2_ balance restoration. Altogether, these observations suggest that the H_2_O_2_ resistance defect in the *zwf1Δ* mutant is, at least in part, a regulated process, rather than merely a consequence of reduced proliferation due to putative cellular damage or a lack of NADPH for anabolic processes.

Similarly, we hypothesized that PKA-driven growth inhibition in the zwf1Δ mutant might also contribute to its H_2_O_2_ tolerance. Using a 4-hour 0.5 mM H_2_O_2_ pulsed stress survival assay, we found that *PDE2* deletion abolished the enhanced survival observed in the *zwf1*Δ mutant (Figure 5G), a result that was further confirmed under higher stress concentrations (Figure S7C and S7D). Moreover, a stress ramping protocol amplified the genetic interaction between *pde2Δ* and *zwf1Δ*: while both single mutants displayed high survival fractions in ramp assays, the vast majority of *pde2*Δ *zwf1*Δ double-mutant cells died, and this effect was exacerbated by supplementing the medium with cAMP (Figure 5H). These results demonstrate that the H_2_O_2_ hyper-tolerance observed in the *zwf1*Δ mutant requires PKA inhibition. Combining the resistance and survival results indicates that the restoration of cell proliferation at low-stress concentrations in the double mutant comes at the expense of significantly reduced survival at higher doses. This observation suggests that PKA regulation plays a decisive role in establishing the balance between resistance and tolerance in response to stress.

How is the internal H_2_O_2_ signal relayed to mediate PKA inhibition in response to stress? Recent studies suggest that oxidized peroxiredoxins - or thioredoxins - may inactivate PKA through a redox-dependent mechanism^40,41^. To further evaluate this model in the context of stress tolerance, we investigated whether deleting thioredoxins or peroxiredoxins would abolish the enhanced survival observed in the *zwf1*Δ mutant. Upon exposure to a 0.5 mM H_2_O_2_ step stress, we found that the high tolerance of both the *zwf1Δ* and *trr1Δ* mutants was suppressed when combined with the *trx1/2*Δ mutations (Figure 5J). Similarly, knocking out all three peroxiredoxins (*TSA1*, *TSA2*, and *AHP1*; referred to as 3PrxΔ) abolished the hyper-tolerant phenotype of the *zwf1Δ* mutant, whereas deletion of *TSA1* alone did not have the same effect (Figure 5J). In line with this, the *trx1/2Δ zwf1Δ* double mutant did not exhibit any fold increase in Msn2-GFP nuclear localization upon H_2_O_2_ exposure, suggesting that PKA was no longer inhibited in this mutant. In contrast, the *tsa1Δ zwf1Δ* mutant maintained Msn2-GFP nuclear localization upon stress (Figure 5K). Hence, these results suggest that the increased tolerance observed in the *zwf1Δ* and *trr1Δ* mutants arises from the activation of a redox signaling relay, rather than being merely a consequence of their reduced growth rate.

Altogether, this genetic analysis suggests that the regulation of PKA activity is a key determinant of cell fate in response to H_2_O_2_ stress, with its modulation shifting the trade-off between resistance and tolerance. To determine whether this trade-off also holds under more physiologically relevant conditions, we next tested the effect of glucose concentration perturbations — known to impact the activation of the Ras-cAMP-PKA pathway — on H_2_O_2_ stress resistance, tolerance, and H_2_O_2_ imbalance (Figure 5L).

To this end, we took advantage of the constant medium replenishment provided by the microfluidic device to grow cells in varying steady glucose concentrations, ranging from 2% to 10_−5_%, which decreased the budding rate from 0.014 min^−1^ to 0.004 min^−1^, respectively (see black curve in Figure 5M). Under these conditions, a mild 0.1 mM H_2_O_2_ stress induced a glucose-dependent decline in the budding rate at steady state: while cells fully adapted to this stress concentration at 2% glucose, their growth rate was permanently reduced at lower glucose concentrations compared to the no-stress condition (see blue curve in Figure 5M). Similarly, upon exposure to an intermediate 0.5 mM H_2_O_2_ stress, reducing the glucose concentration from 2% to 10^−5^% caused a dramatic decrease in the fraction of resistant cells (Figure 5N).

To determine whether this drop in resistance was associated with both a failure to induce the Yap1 regulon and to maintain internal H_2_O_2_ balance, we quantified the response of a strain carrying the Srx1pr-GFP-degron reporter under 0.1 mM H_2_O_2_ stress in cells grown at various glucose concentrations. We found that lower nutrient levels led to a reduction in the reporter’s peak expression (Figure 5O), accompanied by a more pronounced H_2_O_2_ imbalance at steady state, as indicated by a ∼2.5-fold reduction in the HRI from 2% to 10^−5^% glucose (Figure 5P). These results demonstrate that a loss of H_2_O_2_ homeostasis efficacy accompanies the drop in resistance under low-glucose conditions.

Conversely, reducing glucose concentration significantly increased survival in response to a 64 mM H_2_O_2_ stress, ranging from ∼0.05 at 2% glucose to ∼0.6 at glucose concentrations above 10^−3^% (Figure 5Q). Notably, our analysis revealed a “sweet spot” at 0.01% glucose, where cells exhibited both increased resistance (Figure 5N) and tolerance (Figure 5Q) to H_2_O_2_. This effect could be explained by cells maintaining sufficiently high NADPH levels to scavenge H_2_O_2_ while benefiting from reduced PKA signaling, thereby promoting tolerance.

Altogether, these results indicate that glucose concentration dramatically affects cellular fate in response to H_2_O_2_. They provide evidence that the trade-off first observed in the *zwf1Δ* mutant reflects a broader physiological property, essential for understanding how cells modulate growth and survival in response to oxidative stress.

### The trade-off between H_2_O_2_ resistance and tolerance in G6PDH-deficient cells is conserved in the prokaryote *E. coli*

The redox homeostatic system, including both the Prx cycle and the PPP as the main source of NADPH, is highly conserved in other organisms^12,34,55^. In the prokaryote *E. coli*, rerouting of glucose into the PPP in response to H_2_O_2_ is a key metabolic adaptation^49^. We thus thought to explore whether the trade-off between resistance and tolerance revealed by the *zwf1*Δ and *trr1*Δ mutants in *S. cerevisiae* between resistance and tolerance was also conserved in *E. coli*. Mutants deleted for *zwf* (G6PDH, ortholog of *ZWF1* in budding yeast) or *trxB* (thioredoxin reductase, ortholog of *TRR1* in budding yeast) gene had a lower growth rate than the WT (Figure S11A), in agreement with a reduced NADPH level and/or reduced H_2_O_2_ scavenging. We next used a modified mother machine microfluidics device where media flows through the growth channels (Figure 6A, also see Methods) to assess the response to H_2_O_2_ of these mutants (Figure 6B and S11B). Below the MIC (until 0.25 mM for WT and Δ*zwf* and 0.2 mM for Δ*trxB*, Figure S11C), we found that resistant Δ*zwf* cells but not resistant Δ*trxB* cells elicited reduced proliferation (normalized to no stress control) compared to the WT (Figure 6C, also see Methods). Indeed, while the growth rate of WT and Δ*zwf* cells was similarly affected by the treatment at 0.05 mM (see Figure 6D left panel, median normalized growth rate = 0,98 and 0,99 for WT and Δ*zwf* respectively, *p>0.05*), two subpopulations were observed at 0.25 mM, with a significantly higher fraction of cells exhibiting very low growth rate in the mutant (Figure 6D, right panel, median normalized growth rate = 0,71 and 0,32 for WT and Δ*zwf* respectively, *p<0.0001*). Yet, strikingly, the Δ*zwf* mutant (but not Δ*trxB*) had a greater survival fraction than the WT at higher H_2_O_2_ concentrations (Figure 6E), highlighting its increased tolerance. Altogether, these results demonstrate that the trade-off between H_2_O_2_ resistance and tolerance is conserved in the Δ*zwf* but not in the Δ*trxB* mutant in *E. coli*.

**Figure 6:**
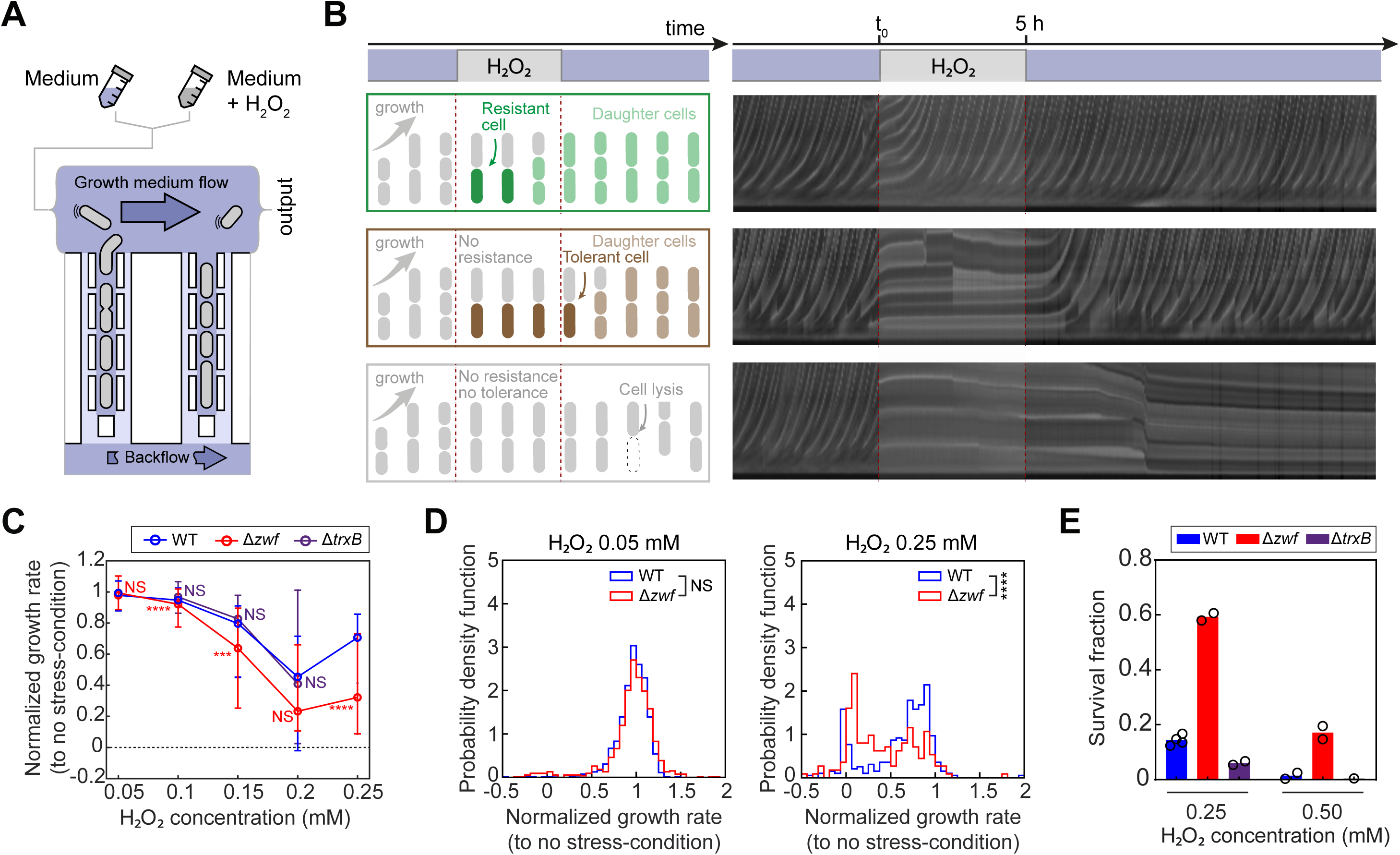
The trade-off between H_2_O_2_ resistance and tolerance in G6PDH-deficient cells is conserved in the prokaryote *E. coli*. **(A)** Sketch of the modified mother-machine microfluidic device used to monitor *E. coli* cells during and after an H_2_O_2_ stress exposure. **(B)** Representative kymographs of three individual cell traps during an exposure to a 0.25mM H_2_O_2_ stress over 5h. Each kymograph represents a particular cell fate (resistance, tolerance, or death). **(C)** Normalized growth rate of cells (relative to the no stress control) as a function of H_2_O_2_ concentration (n>105 for WT, n>185 for Δ*zwf*, and n>90 for ΔtrxB). Points and error bars represent median and interquartile range, respectively. **(D)** Probability density function of the normalized growth rate shown in **(c)** at 0.05 and 0.25 mM H_2_O_2_, in WT and Δ*zwf* strains. **c,d,** Statistical analysis employed one-sided Man Whitney U tests, **** for *p* < 0.0001, *** for *p* < 0.001, ** for *p* < 0.01, * for *p* < 0.05, NS for *p* > 0.05. **(E)** Survival fraction in response to boluses of H_2_O_2_ (0.25 or 0.5 mM for 5 h). Open circles are technical replicates (N = 2 to 4 replicates for WT and Δ*zwf* and Δ*trxB* at 0.25 mM. Only one replicate for ΔtrxB at 0.5 mM since all cells were almost dead at 0.25 mM).

## Discussion

In this study, we sought to uncover the intricate ways in which cells orchestrate diverse defense strategies to ensure their adaptation to challenging environments. To achieve this, we investigated the cellular response to hydrogen peroxide stress in yeast, focusing on the regulation of resistance and tolerance as two distinct yet complementary defense strategies. We developed microfluidics-based assays to independently measure these properties (Figure 1). Our analysis revealed that the Yap1-mediated transcriptional response correlates with resistance but not tolerance (Figure 2). Through a candidate-gene approach, we further showed that mutants associated with the Prx/Trx recycling pathway and NADPH synthesis exhibited decreased resistance, underscoring the importance of redox homeostasis in promoting resistance (Figure 3). Strikingly, we found that deleting *ZWF1* (and its ortholog zwf in E. coli) or *TRR1* revealed an unexpected trade-off with reduced resistance but increased tolerance (Figures 4 and 6). The analysis of genetic interactions suggest that Prx/Trx-dependent redox signaling inhibits PKA, thereby diminishing cell proliferation and maximizing cell survival during stress (Figure 5).

### Diversity of cell fate under H2O2 exposure

Classical H_2_O_2_ “sensitivity” measurement assays using solid media supplemented with hydrogen peroxide cannot discriminate between resistance and tolerance, as both proliferation and survival contribute to the overall cellular fitness and biomass production. This limitation may explain why the hyper-tolerant phenotype of the *trr1*Δ and *zwf1*Δ mutants was not identified in previous studies^12,33,47–50,56^. In contrast, longitudinal cell analyses based on live imaging, as performed in our study, or complementary approaches to score cell growth and survival separately^57^, provide an adequate framework to discriminate among the different possible fates in response to stress. Notably, former studies took advantage of differential stress assays (i.e. growth adaptation on H_2_O_2_ plates and survival to transient H_2_O_2_ stress in liquid cultures) to decipher the specific phenotypes observed by deleting different components of the H_2_O_2_ stress response^37,57^. Interestingly, Fomenko et al. previously identified thioredoxins as crucial for growth adaptation under H_2_O_2_, although their deletion didn’t affect cell survival^57^.

In line with this, our study underlines the non-redundancy of the genes involved in the H_2_O_2_ stress response. In contrast to the NADPH-dependent Prx/Trx pathway which drives H_2_O_2_ scavenging and is the key player in ensuring cellular resistance, cytochrome c peroxidase (*CCP1*), catalase T (*CTT1*) and the general stress response mediated by Msn2/4 specifically contributes to H_2_O_2_ tolerance. Moreover, upon abrupt H_2_O_2_ exposures, cell fate might not only depend on the transcriptional H_2_O_2_ stress response but also on the initial cell state. Consistent with this, previous studies have suggested that pre-conditioning dictates future survival to environmental stressors^45,58^. In our study, survival to severe H_2_O_2_ exposures was indeed correlated with redox imbalance before stress exposure in *zwf1*Δ cells (Figure 4), underscoring the importance of the initial cell state for H_2_O_2_ tolerance.

The use of step and ramp stress patterns was crucial to disentangle these embedded phenotypes. Stress ramping enables cells to reach an “optimal steady-state” by fully activating their transcriptional stress response, while stress stepping preferentially probes the acute capacity of cells to transiently withstand the stressor. Similarly, Kaplan et al. reported that exposing bacteria to stressors either abruptly or gradually results in distinct cell states - disrupted or regulated, respectively^59^. Therefore, modulating stress patterns represents an efficient method to decipher complex cell fate decisions upon stress exposures and may provide valuable insights into other stress response contexts.

### A nutrient-dependent trade-off between stress resistance and tolerance

Several lines of evidence suggest the existence of a trade-off between H_2_O_2_ resistance and tolerance strategies. The increased survival of the *zwf1*Δ and *trr1*Δ mutants compared to WT is associated with a reduction in cell proliferation under stress. This effect is also observed within an isogenic population of *zwf1*Δ cells. Additionally, reducing glucose levels improves the tolerance of WT cells to hydrogen peroxide while decreasing their ability to resist and detoxify H_2_O_2_. Importantly, we can rule out the possibility that the hyper-tolerance observed in the *zwf1*Δ mutant is only due to the lack of NADPH since the level of NADPH is not affected in the *trr1*Δ mutant.

Interestingly, recent studies showed that carbon rerouting into the PPP enhances NADPH production within seconds following H_2_O_2_ exposure^12,33,60^, through allosteric processes such as glycolysis inhibition^56,61^. In PPP deficient cells, H_2_O_2_ exposure thus leads to a rapid drop of NADPH in cells. Therefore, the capacity of cells to sustain the pool of NADPH might dictate stress response strategy within seconds following stress exposure. Extending our study to E. coli revealed the partial conservation of the resistance-tolerance trade-off in the Δ*zwf* mutant (the ortholog of *ZWF1*) but not in Δ*trxB* (ortholog of *TRR1*). While it remains unclear why the trxB mutant did not exhibit the same phenotype as in yeast, we speculate that part of the *zwf* tolerance phenotype might depend on NADPH-dependent but Prx/Trx-independent processes in *E. coli*, such as the antioxidant glutathione^62^ or anabolic processes^50^. Nonetheless, our findings suggest the existence of an evolutionary conserved determinant of H_2_O_2_-defense strategy in eukaryotes and prokaryotes.

Since NADPH is a major metabolite in cells, our work reinforces previous studies highlighting the critical role of resource allocation in stress response, which affects both cellular fitness^63,64^ and stress response^65^. In our study, limiting glucose availability in WT cells induced a hypertolerant phenotype, in line with long-standing observations that starved cells exhibit increased stress survival^19,66,67^. However, we found that reduced glucose availability was also associated with impaired H_2_O_2_ scavenging and resistance, recapitulating phenotypes similar to those observed in the *zwf1*Δ mutant. More broadly, this suggests that starved cells might survive harsh stressors in an ‘*out-of-homeostasis*’ state, relying on protective mechanisms that limit the toxicity of H_2_O_2_, independently of their incapacity to detoxify the stressor. In this context, tolerance appears to function as a last-resort defense strategy, preventing cell death when internal homeostasis is lost, and resistance is no longer functional.

These observations suggest a new perspective on how cells may adjust their stress defense strategies based on resource allocation. We speculate that similar differential strategies could be employed in response to other environmental stressors. For instance, a recent study demonstrated that adaptation to osmolarity is strongly influenced by glucose availability^68^. The coordination between cell resources and stress defense mechanisms might therefore represent a universal feature of stress response that remains to be further investigated.

### A potential PKA/Prx redox relay enables the switch from stress resistance to stress tolerance

Previous studies on the regulation of PKA activation upon H_2_O_2_ exposure have provided genetic and biochemical evidence that its inhibition is mediated either by peroxiredoxins or thioredoxins, which relay H_2_O_2_ signals and physically interact with PKA subunits^40,41^. These findings somewhat downplay the role of peroxiredoxins as the main H_2_O_2_ scavenging enzymes. Our results emphasize that both functions—H_2_O_2_ scavenging and redox signaling—may be equally important: the peroxidase activity is essential for homeostatic system function and resistance, whereas Prx and Trx signaling to PKA could be key mechanisms driving tolerance to hydrogen peroxide (see Figure 7A).

**Figure 7:**
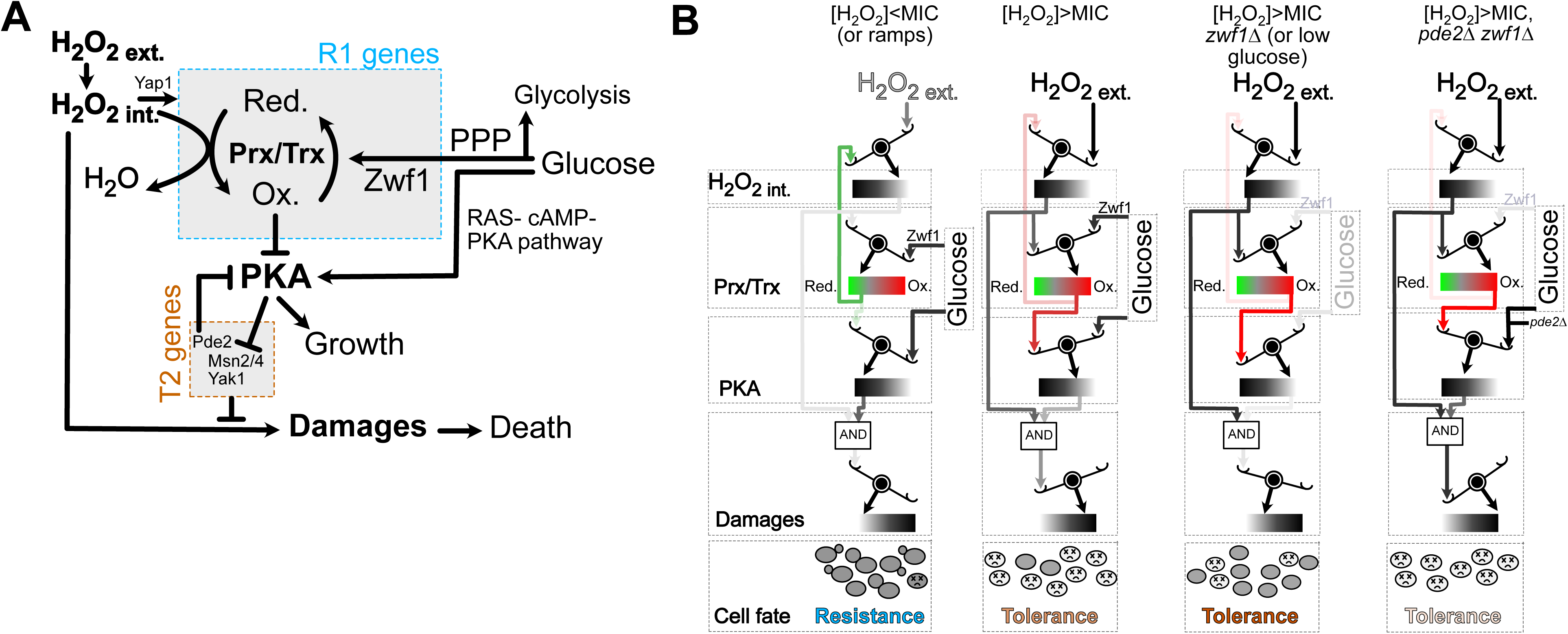
proposed model for the control of cell proliferation and survival in response to hydrogen peroxide. **(A)** Schematic illustrating the interactions between key players described in this study, highlighting resistance genes (**R1** rectangle) and tolerance genes (**R2** rectangle) identified in the analysis, along with their interactions. **(B)** Symbolic model illustrating the system’s functional states in response to varying external H_2_O_2_ levels, genetic mutations, and/or environmental perturbations, along with corresponding cell fates (resistance of tolerance). Each “graded scale” represents the state of key components of the system, including internal H_2_O_2_ levels, the oxidation status of the Prx/Trx machinery, PKA activity, and the level of cellular damage. The opacity of the colored arrows indicates the magnitude of the effect exerted by a given component on a specific target. The “AND” module represents a logical gate, modeling the requirement for both high internal H_2_O_2_ levels and high PKA activity to induce cellular damage (e.g. high PKA activity leads to low expression of tolerance genes and hence mortality only if there is a high level of internal H_2_O_2_).

In this speculative model, a cell’s commitment to a defined H_2_O_2_ response strategy depends on the real-time assessment of the oxidation level of the components of the Prx/Trx pathway, see Figure 7B^41,69^. Under low H_2_O_2_ levels or when H_2_O_2_ levels progressively increase (i.e., ramping stress), the H_2_O_2_ homeostatic system makes necessary adjustments, such as enhancing scavenging capacity through antioxidant production and upregulating NADPH synthesis. In this context, Prx/Trx enzymes remain predominantly reduced, PKA activation supports cell proliferation, and cells avoid accumulating damage that could compromise survival. In contrast, during acute (high) H_2_O_2_ exposure, elevated internal H_2_O_2_ becomes deleterious to cellular function. However, oxidized Prx/Trx enzymes drive PKA inhibition, facilitating the expression of pro-tolerance genes. This limits cell death and ensures a basal level of stress tolerance. In the zwf1Δ mutant or under glucose-limited conditions, the Prx/Trx system is less efficiently reduced, resulting in further PKA inhibition even before stress exposure. This preemptive inhibition enhances the mitigation of damage upon stress exposure, thereby improving tolerance. Conversely, forcing PKA activation in this context prevents the activation of tolerance genes, leading to increased cell mortality. Altogether, this speculative model provides a framework to explain how stress defense strategies can be coordinated to enhance overall cell fitness in fluctuating environments, by restricting growth shutdown to situations where cell survival is critically threatened.

### The interplay between tolerance and resistance beyond H_2_O_2_ stress response

Tolerance is a central mechanism underlying antibiotic treatment failures^6,70,16^ and is associated with specific stress response pathways^66,71^. However, many studies on homeostatic systems primarily focus on understanding the determinants of resistance^4,72–74^. Our analysis of the H_2_O_2_ stress response reveals that resistance and tolerance are intertwined elements with overlapping molecular bases, both contributing to cellular fitness. Notably, mutations that impair resistance, such as *zwf1Δ*, do not necessarily affect tolerance and may even enhance it. Conversely, preventing tolerance by over-activating PKA (e.g., in the *pde2Δ* mutant) does not necessarily impair cellular resistance (pde2Δ mutant has no growth defect in H_2_O_2_ ramps). Importantly, altering both resistance and tolerance results in a significant fitness defect regardless of the temporal stress profile (see Figure 5). Many therapeutic strategies attempt to impair cellular functions by targeting proliferation (i.e., resistance). Yet, powerful tolerance mechanisms can lead to treatment relapse. Therefore, targeting both defense strategies simultaneously could provide a more effective approach to preventing relapse and improving treatment outcomes.

## Methods

### Strains and plasmids

All budding yeast strains were congenic to the S288c background (Sikorski and Hieter, 1989; Huh et al., 2003) and derived from BY4741 or BY4742. The list of strains is detailed in a dedicated supplementary file. Simple mutant strains were all taken from the BY4742 delta collection (invitrogen). The *trx1*Δ *trx2*Δ and the *tsa1*Δ*tsa2*Δ*ahp1*Δ strains were gifts from the Toledano Lab and were also derived from S288c. The strain *msn2*Δ*msn4*Δ was a gift from *Dr.* Li Wei and was also derived from S288c. All strains have been genotyped by PCR. The transcriptional reporter strains carrying the *SRX1pr-sfGFP-deg* were generated by a one-step cloning-free method (Huber et al., 2014) in the corresponding mutant strain issued from the BY4742 delta collection (invitrogen). The double mutants (*X*Δ *zwf1*Δ*::natMX4*) were obtained by substituting the entire *ZWF1* gene by a natMX4 cassette in the corresponding *X*Δ*::kanMX4* strain from the BY4742 delta collection (invitrogen). The mutants *tsa1*Δ*tsa2*Δ*ahp1*Δ*zwf1*Δ*::natMX4, trx1*Δ*trx2*Δ*zwf1*Δ*::natMX4* and *trx1*Δ*trx2*Δ*trr1*Δ*::natMX4* were obtained following the same procedure, substituting the *ZWF1* or the *TRR1* gene by a natMX4 cassette in the corresponding strain. The protein fusion GFP strains were obtained from the BY4741 Invitrogen collection. The strains HSP104-GFP *zwf1*Δ*::natMX4* and SRX1-GFP *zwf1*Δ*::natMX4* were obtained by substituting the entire *ZWF1* gene by a natMX4 cassette in the corresponding protein fusion GFP strain (from invitrogen).

For bacterial strains, the ortholog of yeast thioredoxin reductase trr1 and of zwf1 in *E. coli* are *trxB* and *zwf* respectively. The Δ*trxB* strain was obtained from the KEIO library ^75,76^. The Δ*zwf* strain of the collection was incorrect, and was therefore constructed from scratch. After verification by sequencing, it was moved into BW25113 using generalized P1 transduction. Deletions of *trxB* and *zwf* were confirmed by PCR and subsequent Sanger sequencing (Microsynth) using locus-specific primers. Additionally, we transformed BW25113 with a low-copy plasmid pUA66-hi2GFP bearing synthetic-promoter-driven GFP to monitor the occurrence of lysis (detected as sudden loss of GFP content). We refer to this strain as “WT.”

### Microfabrication and microfluidics setup

The microfluidic chips design for yeast cell experiments was the same as that in Goulev et al.^7,77^. Microfluidics chips were PDMS (Sylgard 184,10:1 mixing ratio) replicas of the master mold. The chip was covalently bonded to a 24 × 50 mm coverslip using a Plasma activator (Diener, Germany). The microfluidic chip was connected to a peristaltic pump (Ismatec, Switzerland) within one hour following the plasma activation (Diener zepto, Germany) with Teflon tubing. The flow rate of the pump was set between 20 and 60 μL/min (see specific protocol for the generation of H_2_O_2_ step and linear ramps below). Hyper7 experiments were performed in a separate microfluidic chip similar to that developed in Aspert et al^78^.

For *E. coli* experiments, we modified the design of the “mother machine” microfluidic device so as to let media flow through the growth channels while retaining cells. To do so, we added shallow structures (0.35*μ*m) around the growth channel and connected those to another flow channel at the back, as described elsewhere^79^. To ensure rapid and controlled switching between conditions, we utilized the same “dial-a-wave” mixer from the DIMM device described earlier ^80^, although without mixing serpentine since we only used one media input or the other; as previously, the chip is designed in order to perform experiments with 8 strains and/or pairs of media in parallel. We manufactured the devices using standard methods of soft lithography. After cutting the chip and punching inlets, the chip was bonded to a pre-cleaned glass coverslip (Schott, TD_00134) by air plasma activation (20-25 s at pressure between 15-20 Pa, power ‘High’ on plasma cleaner PDC-32G, Harrick Plasma) immediately before assembling the chip, followed by baking for 1-1.5 h at 80°C. We assembled the device and primed it with deionized water on the day of the experiment.

### Growth medium and H_2_O_2_ preparation

For yeast experiments, precultures and microfluidic experiments were done at 30°C in a synthetic dextrose (SD) medium, containing a yeast minimal synthetic medium (Takara 630412, yeast nitrogen base, ammonium sulfate and 2% dextrose) supplemented with all amino-acids (Takara 630308 aminoacids dropout mix supplemented with the missing amino-acid). For *E. coli* experiments, precultures and microfluidic experiments were done at 37°C, using filter sterilized M9 minimal medium (Sigma Aldrich, M6030), supplemented with 0.2% w/v glucose (AppliChem, A1349) and 0.1% w/v casamino acids (Sigma Aldrich, 22090). The H_2_O_2_ (Hydrogen peroxide solution 35wt. % in H_2_O_2_, 349887–500 mL, Sigma) stock solutions were prepared and stored as described in Goulev et al., eLife 2017. In brief, for all experiments, H_2_O_2_ was added to the medium just before the experiment to prevent its degradation. We previously showed that in these conditions, the H_2_O_2_ concentration in the tank medium was stable enough for long term microscopy experiments ([H_2_O_2_] decrease was less than 10% in 16,5h) ^7^. We stored the stock solution of hydrogen peroxide at 4°C in the dark after opening.

### Generation of H_2_O_2_ steps and linear H_2_O_2_ ramps

To generate an H_2_O_2_ step pattern, we manually switched the microfluidic device’s medium from synthetic dextrose (SD) medium to an SD medium enriched with H_2_O_2_ at specified concentrations. This switch was executed with precision to avoid introducing air bubbles. Previously, we characterized the diffusion kinetics in this device following a step-like transition to a medium containing fluorescein^7^. The half-rise time for fluorescein diffusion was 21 seconds, significantly faster than the 120-second half-time observed for Yap1 relocalization in the nucleus under H_2_O_2_ stress. Consequently, it is likely that the cells rapidly detect a step-like increase in H_2_O_2_ concentration using this method.

To produce a linear ramp of H_2_O_2_ concentration, we employed a previously described protocol^7,77^. Briefly, a peristaltic pump infused a fresh stock solution of 112.1 mM H_2_O_2_ into the medium tank feeding the microfluidic device. We carefully adjusted the flow rate (µ0) of the pump delivering the H_2_O_2_ stock to match the rate (µ1) at which another pump transferred medium from the tank to the device. This balance maintained a constant volume (V1) in the medium tank over time, ensuring a steady increase in H_2_O_2_ concentration without volume changes (dV1/dt = 0).

In these steady conditions, we can simplify the evolution of C_1_ over time as :

C_1_ = C_0_.(1-e^µ0/V1.t^)

When t << µ_0_/V_1_, we can therefore approximate the evolution of C_1_ as :

C_1_ = C_0_.µ_0_/V_1_.t

With µ_0_ = 50µL/min and V_1_ = 1L, C_1_ follows a linear increase for t << 10^5^ min, therefore until timescales much longer than our experiments (∼240 min of ramp increase).

The tank connected to the microfluidics device was not kept on ice as it required continuous stirring. Additionally, due to the observed degradation of H2O2 in the tank, we calculated a corrective factor from experimental data to adjust the initial concentration of H2O2, denoted as C1. The formula used was:

C_1_ = 0.785.C_0_.µ_0_/V_0_.t

In our study, we controlled the rate of H2O2 increase to 4.4 µM/min. This rate was chosen based on previous findings showing that it allows wild-type (WT) cells to fully activate their transcriptional response. To achieve this, we utilized a constant flow rate of *μ*0=50 µL/min and a tank volume*V*0=1 liter. The initial concentration *C*0 was set at 112.1 mM, enabling the desired gradual increase in H2O2 concentration.

In figure 2, the ramp pattern was designed in such a way that the integrated dose and the maximal dose of [H_2_O_2_] was the same as the step pattern (integrated dose was 0.5 x 240 = 120 mM.min and maximal dose = 0.5 mM). Using a 4.4 µM/min linear increase of H_2_O_2_ for the ramp assay, H_2_O_2_ concentration reached 0.5 mM in about 114 min. After reaching 0.5 mM, the maximal concentration was maintained for 183 min; half of the duration of the ramp increase (57 min) was subtracted from the 240 min so that the integrated dose of both assay was equal. In this assay, stress tolerance was evaluated by the post-stress survival fraction after the 4h stress duration. Stress resistance was evaluated through the quantification of the fold-change in the number of resistant micro-colonies over the 4-hour stress period. For the sake of simplicity, this method of quantification was preferred over the more tedious measurement of the Minimum Inhibitory Concentration (MIC).

### Culture conditions and Time-lapse microscopy

For budding yeast experiments, we streaked yeast strains on YPD agar plates from frozen glycerol stock and let them grow at 30°C for at least two days. Then, one day before timelapse imaging, cells were pre-grown overnight at 30°C in synthetic dextrose (SD) medium from a single colony on the plate (see growth medium section for details). Overnight cultures were then diluted and grown for 4 to 6h to mid-log phase and injected in the microfluidics device, using a 1mL syringe and a 21G needle. Cells were left to grow at least 90 min in the microfluidic chip before the beginning of the experiment. For time-lapse imaging of yeast cells, images were taken using an inverted Zeiss Axio Observer Z1 or a Zeiss Axiovert. The focus was maintained using a software custom algorithm developed on MatLab. Fluorescence images were taken using a LED light (CoolLed, LumenCor) and an EM-CCD Luca-R camera (Andor) and using either a 63x objective (with Zeiss Axio Observer Z1) or a 40x objective (Zeiss Axiovert). Multi-Position imaging was enabled by an automated stage (up to 80 positions). The temperature was set to 30°C during the whole experiment using both a custom objective heater (controlled with a 5C7-195, Oven Industries) and a holder heater (controlled using a custom Arduino module).

For *E. coli* experiments, we streaked bacterial strains on LB agar plates with antibiotics from frozen glycerol stocks and incubated the plates overnight at 37°C before the experiment. On the day of the experiment, we inoculated liquid cultures from freshly grown colonies, and harvested bacteria at OD 0.01-0.05 for loading the microfluidic device. After starting the setup and its temperature control, we let it equilibrate several hours in advance. We used a pressure controller (OB1 mk3+, Elveflow) to control the flows in the microfluidic chip during the experiment. After mounting the chip, we ran the growth medium through for 1-3 h using our standard flow of ≈ 1.6 *μ*L/min per individual series (1.56 − 1.44 × 10^5^ Pa inlet pressure, −10^4^ Pa vacuum). Next, we loaded cells from the flow channel outlet by (i) increasing the flow through growth channels (media inlets at 8 × 10^4^ Pa and overflow channel at −6×10^4^ Pa) and (ii) pressurizing the loading tubing using a manifold connected to the pressure controller. Increasing the pressure on the cell outlet pushed the bacteria into the growth channels. To monitor the switching between different media, we added fluorescein (Sigma Aldrich, F6377) in media containing hydrogen peroxide at a final concentration of 10 ng/mL. An inverted Nikon Ti2-E microscope, equipped with a motorized xy-stage and enclosed in a temperature incubator (TheCube, Life Imaging Systems), was used to perform microfluidic experiments. Images were recorded using a CFI Plan Apochromat Lambda DM 100× oil objective (NA 1.45, WD 0.13 mm) and a sCMOS camera (Photometrics Kinetix); the largest possible field of view of 208 µm×208 µm (3200 pixel×3200 pixel) was obtained by using large optical elements in the light path. The setup was controlled using µManager 2^81^ and time-lapse movies were recorded with its Multidimensional-Acquisition engine. At each position, we acquired every 3 min a phase-contrast image using 110 ms exposure and an image of GFP fluorescence (130 ms exposure, 50% power attenuated by OD 1 filter, Lumencor Spectra 3, Cyan LED with ex 475/35 nm; em 525/50 nm; bs 495 nm filters). We chose these acquisition settings and frequency as they minimize the phototoxicity due to exposure to the short-wavelength excitation light and yet still allow for around ten acquisitions between cell divisions. For each condition-strain combination, we were able to acquire six to seven positions (around 50-60 growth channels per position).

### Image analysis for yeast experiments

For budding yeast experiments, raw images were imported and processed on MatLab using a free access custom software (phyloCell, available on GitHub, Charvin 2017). The software was used to segment and track cells over time based on phase contrast images. Segmentation and tracking were then corrected manually with the phyloCell interface when needed for single-cell analysis. Experiments performed with the Hyper7 probe were analyzed using our Detecdiv software, as previously described^78^. Camera background was systematically subtracted for all fluorescence quantifications.

### Quantification of physiological and fluorescence parameters in budding yeast

#### Minimal inhibitory concentration (MIC)

The minimal duration to inhibit the growth of the population was routinely assessed by the incapacity of cells to recover an exponential growth within 12 h following stress exposure. No WT cells were found to recover growth after 12 h if they hadn’t started recovering before that point. In addition, due to the reduced growth rate in the *zwf1Δ* mutant, we tested in three independent replicates that assessing growth recovery for 24 h instead of 12 h did not change its MIC (data not shown).

#### Post-stress survival fraction

For post-stress survival quantification, only cells present at the beginning of the stress were included in the analysis. Following this procedure, the post-stress survival fraction was assessed independently of the proliferative capacity of cells under stress and therefore independently of stress resistance. The survival fraction was manually measured by determining the fraction of cells (born before stress exposure) able to form at least 2 consecutive buds following stress release. Counting two buds enabled us to exclude cells that remained arrested as G2/M during their first post-stress division due to a DNA-damage checkpoint arrest (Goulev et al., 2017).

#### Minimal duration to kill 99% of the population (MDK99)

The MDK99 was calculated following the procedure described in the ‘*Post-stress survival*’ section. Each independent replicate included at least one hundred cells and typically a few hundred cells. This limits the accuracy of the measurement of the survival fraction. The MDK99 of the population was assessed based on 2 to 4 independent microfluidic experiments for each condition tested.

#### Fold change in cell proliferation (Figure 2)

In the mutant screen in Figure 2, proliferation under stress was assessed by measuring the fold change in cell number during the finite stress period for each micro-colony present at the beginning of the stress period. Importantly, only resistant cells were included in the analysis to avoid any artificial drop in proliferation due to non-growing tolerant or dead cells. Normalized cell proliferation was then calculated by dividing the averaged proliferation obtained in the tested condition to that of the no stress WT condition.

Notably, since all mutants were submitted to a unique dose of H_2_O_2_ (0.5 mM) and stress duration (4h), some mutants exhibited small fractions of resistant cells. This resulted in a small number of resistant micro-colonies included in the proliferation analysis for some mutants, even after including all cells from the 3 experimental replicates. In the step assay, mutants with less than n = 30 micro-colonies quantified for proliferation were *ctt1*Δ (n=6), *srx1*Δ (n=7) and *yak1*Δ (n = 8), *3Prx*Δ (n = 14) and *ccp1*Δ (n = 17). In the ramp assay, mutants with less than n = 30 micro-colonies quantified for proliferation were *ctt1*Δ (n = 25), *trx1/2*Δ (n = 21), *3Prx*Δ (n = 10) and *glr1*Δ (n = 22). However, these small numbers didn’t affect our conclusions since it principally affected the quantification of proliferation of the step assay, which was anyway most specifically designed for assessing survival. Proliferation was best captured with the ramp assay in the sensitive mutants. In addition, the key mutants identified in this screen, *trr1*Δ and *zwf1*Δ, were not affected by this problem (n = 41 and 32 in the step assay and n = 33 and 32 in the ramp assay respectively).

#### Post-stress survival fraction (Figure 2)

In the mutant-screen in Figure 2, the same method as described in the general ‘Post stress survival fraction’ section was used to assess stress survival (i.e. tolerance) in the different mutants. For all mutants screened, N = 3 to 5 technical replicates were used to quantify the mean survival fraction. WT mean survival fraction included N = 13 replicates.

#### Bud to bud frequency (Figure 3 and 4)

The bud to bud frequency was calculated as the inverse of the time measured between the formation of two successive buds. This readout is very similar to the quantification of ‘Fold cell proliferation’ but enables a single-cell approach by quantifying the proliferation of each single-cell under stress (in Figure 3 and 4). To avoid taking into account the acute regime following H_2_O_2_ exposure, we excluded the budding events occurring during the first 5 hours of stress exposure to specifically assess the steady-state resistance of cells. Cells exhibiting a bud-to-bud duration >10h were arbitrarily considered as non-budding cells (frequency <0.0017 min^−1^) and therefore as non-resistant cells.

#### Cell-cycle re-entry following stress exposure (Figure 3)

Cell cycle re-entry was quantified manually, assessing the moment at which each single-cells initiated a newly formed bud after stress release; the cumulative fraction of surviving cells re-entering the cell cycle over time was then plotted (see Fig. 3e, dash line).

#### Quantification of the SRX1pr-GFP-deg

The Srx1pr-GFP-deg signal was quantified as the mean pixel value within the segmented cell area. Peak fluorescence was determined in each cell as the level of fluorescence at t=1h following stress addition after subtracting the initial fluorescence value in the absence of stress.

#### HyPer7-based measurements of cytosolic H_2_O_2_ dynamics

Yeast cells transformed with a p413TEF-HyPer7 plasmid were grown in HC medium with 2% (w/v) glucose as carbon source and lacking histidine for plasmid selection. Two types of assays were performed with the Hyper7 probe. First, we exposed the cells growing in a microlfuidic device similar to that previously described in Aspert et al^78^ to a 0.4 mM step stress and we measured the fluorescence using a dual band filter set (Excitation at 400nm and 470nm, emission at 510nm) during a 5-min interval time lapse experiment. This measurement was compared to the signal obtained using the Srx1pr-GFP-degron filter used throughout the manuscript (Figure S3A and S3B). In a second set of experiments (Figure S3C and S3D), cells were grown at 30°C with shaking until the culture reached an optical density of OD_600_ ≈ 3.3. Cells were harvested by centrifugation and resuspended to an OD_600_ = 7.5 in 100 mM MES/Tris pH 6 buffer. Cells were transferred in 200 µL aliquots to a flat-bottomed 96-well imaging plate. HyPer7 probe fluorescence was subsequently monitored using a BMG Labtech CLARIOstar fluorescence plate reader with excitation of 400 and 480 nm and an emission of 510 nm in both cases. Fluorescence was measured for 5 mins before the addition of H_2_O_2_ at 0.2 mM.

#### H_2_O_2_ Restoration Index (HRI)

The H_2_O_2_ restoration index (HRI) was calculated using the Srx1pr-GFP-deg quantification, as the ratio of the mean GFP value 5h after the addition of a 0.1 mM H_2_O_2_ stress, and divided by the mean GFP value 1h after stress addition:

HRI(Srx1pr-GFP-deg) = Fluo(t=1h) / Fluo(t=5h)

This score was then normalized to the WT. A small HRI thus represents a low ability to restore H_2_O_2_ balance. The H_2_O_2_ concentration was chosen to ensure that every strain included in the analysis could grow and display a transcriptional response upon stress exposure, since the signal of non-responder cells might not be interpreted to assess H_2_O_2_ homeostasis recovery.

The H_2_O_2_ restoration index using the ratiometric Hyper7 probe was quantified with a similar procedure, but using different time points to calculate the decay from the oxidized version to the reduced version of the probe following H_2_O_2_ exposure (quantified as the fluorescence ratio 488nm/405nm). Based on the recovery dynamics of the Hyper7 probe in the WT strain following H_2_O_2_ exposure, we defined:

HRI(Hyper7) = Hyper7(t=5min) / Hyper7(t=35min)

#### Yap1 and Msn2 nuclear localization

Yap1-sfGFP and Msn2-GFP nuclear localization were then quantified as described in Cai et al., Nature, 2008. Briefly, the nuclear localization score was measured by the difference between the mean intensity of the 5 brightest pixels in the segmented cell and the mean intensity of all other pixels in the same segmented cell.

#### Quantification of Hsp104-GFP and Tsa1-GFP aggregates

The relative aggregation-scores of fusion protein were quantified using the same methodology as for Yap1 and Msn2 nuclear localization scores, based on the mean intensity of the 5 brightest pixels in the segmented cells as previously described in S. Saad et al., Nat Cell Biol, 2017. We checked that the results were independent of the number of pixels used to assess the specific signal of the GFP foci. To assess the Hsp104-GFP aggregation score following stress release in the *zwf1*Δ mutant (Fig. 3D and E), we excluded dead cells based on their mean fluorescence signal to only account for the Hsp104 deaggregation of viable cells. Due to the very high GFP levels in viable cells, we excluded cells with a mean fluorescence level <1000 a.u. at the moment of stress release. The fluorescence level in viable cells was far above this threshold under H_2_O_2_ (see Extended Data Fig. 3H).

#### Normalized tolerance, resistance and stress response capacities

To compare how H_2_O_2_ tolerance, resistance and stress response behaves as a function of H_2_O_2_ concentration, we defined dimensionless normalized variables for tolerance and stress response. Normalized tolerance was defined as the MDK99 at a given concentration divided by the MDK99 at 1 mM (i.e. the MDK99 right above the MIC, where tolerance starts to be defined). The normalized stress response at t=1h following stress addition was defined based on the mean Srx1pr-GFP-degron expression (see ‘*Quantification of the SRX1pr-GFP-deg’* section) as follows: *Normalized stress response = 1 - ((Exp - Fit) / Fit)*, where *Exp* is the experimentally measured peak expression for each [H_2_O_2_] and *Fit* the extrapolation of the linear fit at small doses of H_2_O_2_, considering a linear dose-response peak expression. Normalized resistance and stress response were then fitted using a sigmoidal function, with a half capacity obtained at very similar H_2_O_2_ concentrations (0.27 and 0.28 mM respectively). Tolerance was not following an exponential decay and was thus fitted using a power law.

#### Prediction of H2O2 stress survival from pre-stress fluorescence (Figure 3)

To evaluate how redox imbalance affects future stress exposure survival, we measured the level of the Srx1-GFP fusion proteins at the onset of a burst of 7.5 or 15 min of 64 mM H_2_O_2_ exposure. The mean normalized fluorescence level was used as a proxy of the redox status of cells before stress. To avoid that experiment-to-experiment variability in the fluorescence level introduced a bias in the prediction, we normalized the fluorescence by the median fluorescence signal for each experiment independently, before pooling cells from all experiments together. Cells were then sorted into 6 groups based on their normalized fluorescent value. The viable/death status of each single cell was then determined as described in the ‘*Post-stress survival fraction*’ section and the average fraction of surviving cells in each group was plotted as a function of the normalized Srx1-GFP fluorescence level prior stress exposure. For these experiments, the GFP strains used were in the BY4741 background; we checked that the tolerance phenotype was not affected when compared to the equivalent strains in the BY4741 background (Figure S1E and S1F).

### Image analysis for *E. coli* experiments

We analyzed the micrographs time-lapses in two stages. First, in the preprocessing step, we isolated individual growth channels, and corrected for small drift experienced during the acquisition. This step created videos of individual channels as well as longitudinal kymographs of growth channels. We used the latter for manually determining the regrowth, and the former for automated image analysis. We segmented and tracked the lineage of bacterial cells until the end of the hydrogen peroxide treatment using DeepMoMA. This software builds on the foundation of the original Mother-Machine analyzer (MoMA) and will be described elsewhere. DeepMoMA utilizes convolutional neural networks for generating cell-outline hypotheses and treats cell segmentation and tracking of the cells as a joint problem. Upon solving the tracking-segmentation problem, cell statistics were exported directly (e.g., cell length, lineage, etc.) without manual curation. The software is available at https://github.com/nimwegenLab/moma/releases/tag/v0.9.6 and we used segmentation model “model_20230509_8616f16b.zip” available at https://github.com/nimwegenLab/moma-model. We estimated the growth rate before the treatment for cells observed during their whole cell cycle. We fitted a linear model to the log-transformed length time-trace vs time. We did not observe any systematic effects in unperturbed growth rate between the different locations in the microfluidic device. Day-to-day variability did not exceed the variability we observed within a single experiment, except in the case of Δ*trxB* where we observed a detectable shift in mean growth rate between replicates. To obtain the fraction of resistant cells, we identified cells that were present at the start of the oxidative stress. A cell was considered non-growing if its cumulative length did not increase by more than 1.8-fold during the treatment. In order to successfully detect the growth status for as many cells as possible, we summed the length of its two daughters if a cell divided during the treatment, and cell traces were considered until the elongation threshold was exceeded, or one of the two daughters was lost (because it left the channel). In order to quantify the reduction of growth rate during oxidative stress, we considered all cells which were observed during at least 4 consecutive time points. In order to focus only on resistant cells, we kept only cells which were labeled as resistant based on the criteria described above. We fitted a linear model to the log-transformed length time-trace vs time to estimate the growth rate, and normalized it to the mean growth rate observed before treatment for the same strain. Exposure to high levels of oxidative stress incurred cell death and lysis. While the cell debris made segmentation of cell outlines difficult, we could estimate the fraction of post-stress regrowth and thus tolerance from kymographs of the growth channels generated during preprocessing. Aided by custom Python scripts, we manually annotated the kymographs for regrowth during and after treatment.

### Bulk growth-rate determination for *E. coli*

We estimated the bulk growth rates by growing cultures in microtiter plates. We grew two different clones of WT, Δ*zwf*, and Δ*trxB* overnight in M9 supplemented with 0.2% glucose, 0.1% casamino acids, and 50 µg/mL kanamycin. The next day, we prepared a plate with the same media, but without kanamycin and with 0.001% TritonX. The addition of TritonX ensured a flat meniscus of the liquid in a well and allowed for a better absorbance measurement. We diluted the overnight cultures around 1:2.6×10^5^, which ensured that the cultures were in steady exponential growth when detected. We acquired data for 16 technical replicates of every clone. We measured the absorbance at 600 nm every 10 min over 24 h with continuous shaking (double orbital, 1 mm kick, 807 cpm) at 37°C using Biotek Synergy H1. This procedure provided 145 reads per growth curve for a given well. We subtracted the absorbance background in a per-well fashion; for a given well, we subtracted the median of the first 5 time samples. For such background-corrected curves, we selected intervals containing points that were at least two doublings away from the stationary phase but did not exceed absorbance of 0.1 nor were below 0.01. We log-transformed these absorbance readings and fitted a linear function, whose slope is the growth rate.

## Author Contributions

BJ: Conceptualization, Methodology, Software, Validation, Formal analysis, Investigation, Writing - original draft, Funding acquisition; BK: Conceptualization, Methodology, Software, Validation, Formal analysis, Investigation, Writing - review and editing; TA: Methodology, Software, Resources; AM: Methodology, Resources; AK: Investigation; AZ: Investigation, Formal analysis; EB: Investigation, Formal analysis; BM: Resources, Formal analysis, Supervision, Funding acquisition; TJ: Conceptualization, Methodology, Software, Validation, Formal analysis, Investigation, Writing - review and editing, Supervision, Funding acquisition; GC: Conceptualization, Methodology, Software, Validation, Formal analysis, Writing - original draft, Supervision, Project administration, Funding acquisition.

## Acknowledgments

We thank Michel Toledano for fruitful discussions, Sophie Quintin, Sandrine Morlot for careful reading of the manuscript. We thank Denis Fumagalli for media and plate preparations. BJ thanks Erwan Grandgirard and Didier Hentsch for their support with microscopy. KB acknowledges tremendous support from M. Mell with the use of deepMoMA and thanks Maria-Elisenda Alaball Pujol for help with microfluidics. KB and TJ are grateful to Erik van Nimwegen and members of his research group for generous support and discussions, which rendered this work possible. Computations carried out by KB and TJ were partially performed at sciCORE (http://scicore.unibas.ch/) scientific computing core facility at University of Basel. This work was supported by the Fondation pour la Recherche Médicale (FRM, B.J, and G.C.), the Agence Nationale pour la Recherche (G.C.), the grant ANR-10-LABX-0030-INRT, a French State fund managed by the Agence Nationale de la Recherche under the frame program Investissements d’Avenir ANR-10-IDEX-0002-02. BM gratefully acknowledges funding from the Deutsche Forschungsgemeinschaft, grant MO2774/6-1; project number, 505680640. KB gratefully acknowledges support by a postdoctoral fellowship of the “Peter und Traudl Engelhorn Stiftung.” TJ acknowledges funding from the Swiss National Science Foundation (SNSF project 310030_197836). BJ gratefully acknowledges funding from an MDPhD student fellowship from Edilb. This project has benefited from establishing collaboration at KITP workshop “Quantitative biology of non-growing microbes” supported by NSF Grant No. PHY-1748958 and the Gordon and Betty Moore Foundation Grant No. 2919.02.

## Declaration of generative AI and AI-assisted technologies in the writing process

During the preparation of this work, the author(s) utilized OpenAI’s ChatGPT-4 to enhance the readability and clarity of the text, given that the primary authors are non-native English speakers. The author(s) subsequently reviewed and edited the AI-generated content as necessary and take full responsibility for the final content of the publication.

## References

1. Veal, E.A., Day, A.M., and Morgan, B.A. (2007). Hydrogen peroxide sensing and signaling. Mol. Cell 26, 1–14.

2. Toledano, M.B., Delaunay, A., Monceau, L., and Tacnet, F. (2004). Microbial H2O2 sensors as archetypical redox signaling modules. Trends Biochem. Sci. 29, 351–357.

3. Toledano, M.B., Planson, A.-G., and Delaunay-Moisan, A. (2010). Reining in H(2)O(2) for safe signaling. Cell 140, 454–456.

4. Hohmann, S. (2002). Osmotic stress signaling and osmoadaptation in yeasts. Microbiol. Mol. Biol. Rev. 66, 300–372.

5. Brauner, A., Fridman, O., Gefen, O., and Balaban, N.Q. (2016). Distinguishing between resistance, tolerance and persistence to antibiotic treatment. Nat. Rev. Microbiol. 14, 320–330.

6. Balaban, N.Q., Helaine, S., Lewis, K., Ackermann, M., Aldridge, B., Andersson, D.I., Brynildsen, M.P., Bumann, D., Camilli, A., Collins, J.J., et al. (2019). Definitions and guidelines for research on antibiotic persistence. Nat. Rev. Microbiol. 17, 441–448.

7. Goulev, Y., Morlot, S., Matifas, A., Huang, B., Molin, M., Toledano, M.B., and Charvin, G. (2017). Nonlinear feedback drives homeostatic plasticity in H2O2 stress response. Elife 6. 10.7554/eLife.23971.

8. Muzzey, D., Gómez-Uribe, C.A., Mettetal, J.T., and van Oudenaarden, A. (2009). A systems-level analysis of perfect adaptation in yeast osmoregulation. Cell 138, 160–171.

9. Milias-Argeitis, A., Summers, S., Stewart-Ornstein, J., Zuleta, I., Pincus, D., El-Samad, H., Khammash, M., and Lygeros, J. (2011). In silico feedback for in vivo regulation of a gene expression circuit. Nat. Biotechnol. 29, 1114–1116.

10. Young, J.W., Locke, J.C.W., and Elowitz, M.B. (2013). Rate of environmental change determines stress response specificity. Proc. Natl. Acad. Sci. U. S. A. 110, 4140–4145.

11. Davies, K.J.A. (2016). Adaptive homeostasis. Mol. Aspects Med. 49, 1–7.

12. Ralser, M., Wamelink, M.M., Kowald, A., Gerisch, B., Heeren, G., Struys, E.A., Klipp, E., Jakobs, C., Breitenbach, M., Lehrach, H., et al. (2007). Dynamic rerouting of the carbohydrate flux is key to counteracting oxidative stress. J. Biol. 6, 10.

13. Ralser, M., Wamelink, M.M.C., Latkolik, S., Jansen, E.E.W., Lehrach, H., and Jakobs, C. (2009). Metabolic reconfiguration precedes transcriptional regulation in the antioxidant response. Nat. Biotechnol. 27, 604–605.

14. Albert, B., Kos-Braun, I.C., Henras, A.K., Dez, C., Rueda, M.P., Zhang, X., Gadal, O., Kos, M., and Shore, D. (2019). A ribosome assembly stress response regulates transcription to maintain proteome homeostasis. Elife 8. 10.7554/eLife.45002.

15. Smith, A., Ward, M.P., and Garrett, S. (1998). Yeast PKA represses Msn2p/Msn4p-dependent gene expression to regulate growth, stress response and glycogen accumulation. EMBO J. 17, 3556–3564.

16. Levin-Reisman, I., Ronin, I., Gefen, O., Braniss, I., Shoresh, N., and Balaban, N.Q. (2017). Antibiotic tolerance facilitates the evolution of resistance. Science 355, 826–830.

17. Gray, J.V., Petsko, G.A., Johnston, G.C., Ringe, D., Singer, R.A., and Werner-Washburne, M. (2004). “Sleeping Beauty”: Quiescence in Saccharomyces cerevisiae. Microbiol. Mol. Biol. Rev. 68, 187–206.

18. Bigger, J. (1944). TREATMENT OF STAPHYLOCOCCAL INFECTIONS WITH PENICILLIN BY INTERMITTENT STERILISATION. Lancet 244, 497–500.

19. Balaban, N.Q., Merrin, J., Chait, R., Kowalik, L., and Leibler, S. (2004). Bacterial persistence as a phenotypic switch. Science 305, 1622–1625.

20. Slatkin, M. (1974). Hedging one’s evolutionary bets. Nature 250, 704–705.

21. Levy, S.F., Ziv, N., and Siegal, M.L. (2012). Bet Hedging in Yeast by Heterogeneous, Age-Correlated Expression of a Stress Protectant. PLoS Biol. 10, e1001325.

22. Lee, J., Godon, C., Lagniel, G., Spector, D., Garin, J., Labarre, J., and Toledano, M.B. (1999). Yap1 and Skn7 control two specialized oxidative stress response regulons in yeast. J. Biol. Chem. 274, 16040–16046.

23. Delaunay, A., Isnard, A.D., and Toledano, M.B. (2000). H2O2 sensing through oxidation of the Yap1 transcription factor. Sci. STKE 19, 5157.

24. Kuge, S., and Jones, N. (1994). YAP1 dependent activation of TRX2 is essential for the response of Saccharomyces cerevisiae to oxidative stress by hydroperoxides. EMBO J. 13, 655–664.

25. Godon, C., Lagniel, G., Lee, J., Buhler, J.M., Kieffer, S., Perrot, M., Boucherie, H., Toledano, M.B., and Labarre, J. (1998). The H2O2 stimulon in Saccharomyces cerevisiae. J. Biol. Chem. 273, 22480–22489.

26. Gasch, A.P., Spellman, P.T., Kao, C.M., Carmel-Harel, O., Eisen, M.B., Storz, G., Botstein, D., and Brown, P.O. (2000). Genomic expression programs in the response of yeast cells to environmental changes. Sci. STKE 11, 4241.

27. Jiang, H., and English, A.M. (2006). Phenotypic analysis of the ccp1Δ and ccp1Δ-ccp1W191F mutant strains of Saccharomyces cerevisiae indicates that cytochrome c peroxidase functions in oxidative-stress signaling. J. Inorg. Biochem. 100, 1996–2008.

28. Chae, H.Z., Chung, S.J., and Rhee, S.G. (1994). Thioredoxin-dependent peroxide reductase from yeast. J. Biol. Chem. 269, 27670–27678.

29. Pedrajas, J.R., Miranda-Vizuete, A., Javanmardy, N., Gustafsson, J.A., and Spyrou, G. (2000). Mitochondria of Saccharomyces cerevisiae contain one-conserved cysteine type peroxiredoxin with thioredoxin peroxidase activity. J. Biol. Chem. 275, 16296–16301.

30. Wong, C.-M., Zhou, Y., Ng, R.W.M., Kung Hf, H.-F., and Jin, D.-Y. (2002). Cooperation of yeast peroxiredoxins Tsa1p and Tsa2p in the cellular defense against oxidative and nitrosative stress. J. Biol. Chem. 277, 5385–5394.

31. Iraqui, I., Kienda, G., Soeur, J., Faye, G., Baldacci, G., Kolodner, R.D., and Huang, M.-E. (2009). Peroxiredoxin Tsa1 is the key peroxidase suppressing genome instability and protecting against cell death in Saccharomyces cerevisiae. PLoS Genet. 5, e1000524.

32. Ralser, M., Wamelink, M.M., Kowald, A., Gerisch, B., Heeren, G., Struys, E.A., Klipp, E., Jakobs, C., Breitenbach, M., Lehrach, H., et al. (2007). Dynamic rerouting of the carbohydrate flux is key to counteracting oxidative stress. J. Biol. 6, 10.

33. Kuehne, A., Emmert, H., Soehle, J., Winnefeld, M., Fischer, F., Wenck, H., Gallinat, S., Terstegen, L., Lucius, R., Hildebrand, J., et al. (2015). Acute Activation of Oxidative Pentose Phosphate Pathway as First-Line Response to Oxidative Stress in Human Skin Cells. Mol. Cell 59, 359–371.

34. Hall, A., Karplus, P.A., and Poole, L.B. (2009). Typical 2-Cys peroxiredoxins--structures, mechanisms and functions. FEBS J. 276, 2469–2477.

35. Broach, J.R. (2012). Nutritional control of growth and development in yeast. Genetics 192, 73–105.

36. Martínez-Pastor, M.T., Marchler, G., Schüller, C., Marchler-Bauer, A., Ruis, H., and Estruch, F. (1996). The Saccharomyces cerevisiae zinc finger proteins Msn2p and Msn4p are required for transcriptional induction through the stress response element (STRE). EMBO J. 15, 2227–2235.

37. Hasan, R., Leroy, C., Isnard, A.-D., Labarre, J., Boy-Marcotte, E., and Toledano, M.B. (2002). The control of the yeast H2O2 response by the Msn2/4 transcription factors. Mol. Microbiol. 45, 233–241.

38. Boisnard, S., Lagniel, G., Garmendia-Torres, C., Molin, M., Boy-Marcotte, E., Jacquet, M., Toledano, M.B., Labarre, J., and Chédin, S. (2009). H2O2 activates the nuclear localization of Msn2 and Maf1 through thioredoxins in Saccharomyces cerevisiae. Eukaryot. Cell 8, 1429–1438.

39. Molin, M., Yang, J., Hanzén, S., Toledano, M.B., Labarre, J., and Nyström, T. (2011). Life span extension and H(2)O(2) resistance elicited by caloric restriction require the peroxiredoxin Tsa1 in Saccharomyces cerevisiae. Mol. Cell 43, 823–833.

40. Bodvard, K., Peeters, K., Roger, F., Romanov, N., Igbaria, A., Welkenhuysen, N., Palais, G., Reiter, W., Toledano, M.B., Käll, M., et al. (2017). Light-sensing via hydrogen peroxide and a peroxiredoxin. Nat. Commun. 8, 14791.

41. Roger, F., Picazo, C., Reiter, W., Libiad, M., Asami, C., Hanzén, S., Gao, C., Lagniel, G., Welkenhuysen, N., Labarre, J., et al. (2020). Peroxiredoxin promotes longevity and H2O2-resistance in yeast through redox-modulation of protein kinase A. Elife 9. 10.7554/eLife.60346.

42. Fourquet, S., Huang, M.-E., D’Autreaux, B., and Toledano, M.B. (2008). The dual functions of thiol-based peroxidases in H2O2 scavenging and signaling. Antioxid. Redox Signal. 10, 1565–1576.

43. Minard, K.I., Jennings, G.T., Loftus, T.M., Xuan, D., and McAlister-Henn, L. (1998). Sources of NADPH and expression of mammalian NADP+-specific isocitrate dehydrogenases in Saccharomyces cerevisiae. J. Biol. Chem. 273, 31486–31493.

44. Lee, P., Cho, B.-R., Joo, H.-S., and Hahn, J.-S. (2008). Yeast Yak1 kinase, a bridge between PKA and stress-responsive transcription factors, Hsf1 and Msn2/Msn4. Mol. Microbiol. 70, 882–895.

45. Berry, D.B., and Gasch, A.P. (2008). Stress-activated genomic expression changes serve a preparative role for impending stress in yeast. Mol. Biol. Cell 19, 4580–4587.

46. Pak, V.V., Ezeriņa, D., Lyublinskaya, O.G., Pedre, B., Tyurin-Kuzmin, P.A., Mishina, N.M., Thauvin, M., Young, D., Wahni, K., Martínez Gache, S.A., et al. (2020). Ultrasensitive Genetically Encoded Indicator for Hydrogen Peroxide Identifies Roles for the Oxidant in Cell Migration and Mitochondrial Function. Cell Metab. 31, 642–653.e6.

47. Pandolfi, P.P., Sonati, F., Rivi, R., Mason, P., Grosveld, F., and Luzzatto, L. (1995). Targeted disruption of the housekeeping gene encoding glucose 6-phosphate dehydrogenase (G6PD): G6PD is dispensable for pentose synthesis but essential for defense against oxidative stress. EMBO J. 14, 5209–5215.

48. Izawa, S., Maeda, K., Miki, T., Mano, J., Inoue, Y., and Kimura, A. (1998). Importance of glucose-6-phosphate dehydrogenase in the adaptive response to hydrogen peroxide in Saccharomyces cerevisiae. Biochem. J 330 (Pt 2), 811–817.

49. Christodoulou, D., Link, H., Fuhrer, T., Kochanowski, K., Gerosa, L., and Sauer, U. (2018). Reserve Flux Capacity in the Pentose Phosphate Pathway Enables Escherichia coli’s Rapid Response to Oxidative Stress. Cell Syst 6, 569–578.e7.

50. Chen, L., Zhang, Z., Hoshino, A., Zheng, H.D., Morley, M., Arany, Z., and Rabinowitz, J.D. (2019). NADPH production by the oxidative pentose-phosphate pathway supports folate metabolism. Nature Metabolism 1, 404–415.

51. Jang, H.H., Lee, K.O., Chi, Y.H., Jung, B.G., Park, S.K., Park, J.H., Lee, J.R., Lee, S.S., Moon, J.C., Yun, J.W., et al. (2004). Two enzymes in one; two yeast peroxiredoxins display oxidative stress-dependent switching from a peroxidase to a molecular chaperone function. Cell 117, 625–635.

52. Hanzén, S., Vielfort, K., Yang, J., Roger, F., Andersson, V., Zamarbide-Forés, S., Andersson, R., Malm, L., Palais, G., Biteau, B., et al. (2016). Lifespan Control by Redox-Dependent Recruitment of Chaperones to Misfolded Proteins. Cell 166, 140–151.

53. Erjavec, N., Larsson, L., Grantham, J., and Nyström, T. (2007). Accelerated aging and failure to segregate damaged proteins in Sir2 mutants can be suppressed by overproducing the protein aggregation-remodeling factor Hsp104p. Genes Dev. 21, 2410–2421.

54. Görner, W., Durchschlag, E., Martinez-Pastor, M.T., Estruch, F., Ammerer, G., Hamilton, B., Ruis, H., and Schüller, C. (1998). Nuclear localization of the C2H2 zinc finger protein Msn2p is regulated by stress and protein kinase A activity. Genes Dev. 12, 586–597.

55. Chae, H.Z., Robison, K., Poole, L.B., Church, G., Storz, G., and Rhee, S.G. (1994). Cloning and sequencing of thiol-specific antioxidant from mammalian brain: alkyl hydroperoxide reductase and thiol-specific antioxidant define a large family of antioxidant enzymes. Proc. Natl. Acad. Sci. U. S. A. 91, 7017–7021.

56. Talwar, D., Miller, C.G., Grossmann, J., Szyrwiel, L., Schwecke, T., Demichev, V., Mikecin Drazic, A.-M., Mayakonda, A., Lutsik, P., Veith, C., et al. (2023). The GAPDH redox switch safeguards reductive capacity and enables survival of stressed tumour cells. Nat Metab 5, 660–676.

57. Fomenko, D.E., Koc, A., Agisheva, N., Jacobsen, M., Kaya, A., Malinouski, M., Rutherford, J.C., Siu, K.-L., Jin, D.-Y., Winge, D.R., et al. (2011). Thiol peroxidases mediate specific genome-wide regulation of gene expression in response to hydrogen peroxide. Proc. Natl. Acad. Sci. U. S. A. 108, 2729–2734.

58. Mitchell, A., Romano, G.H., Groisman, B., Yona, A., Dekel, E., Kupiec, M., Dahan, O., and Pilpel, Y. (2009). Adaptive prediction of environmental changes by microorganisms. Nature 460, 220–224.

59. Kaplan, Y., Reich, S., Oster, E., Maoz, S., Levin-Reisman, I., Ronin, I., Gefen, O., Agam, O., and Balaban, N.Q. (2021). Observation of universal ageing dynamics in antibiotic persistence. Nature 600, 290–294.

60. Dick, T.P., and Ralser, M. (2015). Metabolic Remodeling in Times of Stress: Who Shoots Faster than His Shadow? Mol. Cell 59, 519–521.

61. Hurbain, J., Thommen, Q., Anquez, F., and Pfeuty, B. (2022). Quantitative modeling of pentose phosphate pathway response to oxidative stress reveals a cooperative regulatory strategy. iScience 25, 104681.

62. Fahey, R.C., Brown, W.C., Adams, W.B., and Worsham, M.B. (1978). Occurrence of glutathione in bacteria. J. Bacteriol. 133, 1126–1129.

63. Mitchell, A., and Pilpel, Y. (2011). A mathematical model for adaptive prediction of environmental changes by microorganisms. Proc. Natl. Acad. Sci. U. S. A. 108, 7271–7276.

64. Weiße, A.Y., Oyarzún, D.A., Danos, V., and Swain, P.S. (2015). Mechanistic links between cellular trade-offs, gene expression, and growth. Proc. Natl. Acad. Sci. U. S. A. 112, E1038–E1047.

65. Balakrishnan, R., de Silva, R.T., Hwa, T., and Cremer, J. (2021). Suboptimal resource allocation in changing environments constrains response and growth in bacteria. Mol. Syst. Biol. 17, e10597.

66. Radzikowski, J.L., Vedelaar, S., Siegel, D., Ortega, Á.D., Schmidt, A., and Heinemann, M. (2016). Bacterial persistence is an active σ S stress response to metabolic flux limitation. Preprint, 10.15252/msb.20166998 10.15252/msb.20166998.

67. Wood, T.K., Knabel, S.J., and Kwan, B.W. (2013). Bacterial persister cell formation and dormancy. Appl. Environ. Microbiol. 79, 7116–7121.

68. Duveau, F., Cordier, C., Chiron, L., Le Bec, M., Pouzet, S., Séguin, J., Llamosi, A., Sorre, B., Di Meglio, J.-M., and Hersen, P. (2024). Yeast cell responses and survival during periodic osmotic stress are controlled by glucose availability. Elife 12. 10.7554/eLife.88750.

69. Stöcker, S., Maurer, M., Ruppert, T., and Dick, T.P. (2018). A role for 2-Cys peroxiredoxins in facilitating cytosolic protein thiol oxidation. Nat. Chem. Biol. 14, 148–155.

70. Boeck, L. (2023). Antibiotic tolerance: targeting bacterial survival. Curr. Opin. Microbiol. 74, 102328.

71. Harms, A., Maisonneuve, E., and Gerdes, K. (2016). Mechanisms of bacterial persistence during stress and antibiotic exposure. Science 354. 10.1126/science.aaf4268.

72. Mettetal, J.T., Muzzey, D., Gomez-Uribe, C., and van Oudenaarden, A. (2008). The Frequency Dependence of Osmo-Adaptation in Saccharomyces cerevisiae. Preprint, 10.1126/science.1151582 10.1126/science.1151582.

73. Muzzey, D., Gómez-Uribe, C.A., Mettetal, J.T., and van Oudenaarden, A. (2009). A Systems-Level Analysis of Perfect Adaptation in Yeast Osmoregulation. Preprint, 10.1016/j.cell.2009.04.047 10.1016/j.cell.2009.04.047.

74. Mitchell, A., Wei, P., and Lim, W.A. (2015). Oscillatory stress stimulation uncovers an Achilles’ heel of the yeast MAPK signaling network. Science 350, 1379–1383.

75. Baba, T., Ara, T., Hasegawa, M., Takai, Y., Okumura, Y., Baba, M., Datsenko, K.A., Tomita, M., Wanner, B.L., and Mori, H. (2006). Construction of Escherichia coli K-12 in-frame, single-gene knockout mutants: the Keio collection. Mol. Syst. Biol. 2, 2006.0008.

76. Datsenko, K.A., and Wanner, B.L. (2000). One-step inactivation of chromosomal genes in *Escherichia coli* K-12 using PCR products. Proc. Natl. Acad. Sci. U. S. A. 97, 6640–6645.

77. Goulev, Y., Matifas, A., and Charvin, G. (2018). Controllable stress patterns over multi-generation timescale in microfluidic devices. Methods Cell Biol. 147, 29–40.

78. Aspert, T., Hentsch, D., and Charvin, G. (2022). DetecDiv, a generalist deep-learning platform for automated cell division tracking and survival analysis. Elife 11. 10.7554/eLife.79519.

79. Julou, T., Gervais, T., and van Nimwegen, E. (2022). Growth rate controls the sensitivity of gene regulatory circuits. bioRxiv, 2022.04.03.486858. 10.1101/2022.04.03.486858.

80. Kaiser, M., Jug, F., Julou, T., Deshpande, S., Pfohl, T., Silander, O.K., Myers, G., and van Nimwegen, E. (2018). Monitoring single-cell gene regulation under dynamically controllable conditions with integrated microfluidics and software. Nat. Commun. 9, 212.

81. Edelstein, A., Amodaj, N., Hoover, K., Vale, R., and Stuurman, N. (2010). Computer control of microscopes using µManager. Curr. Protoc. Mol. Biol. Chapter 14, Unit14.20.

