## Supplementary material for "A trade-off between stress resistance and tolerance underlies the adaptive response to hydrogen peroxide": Suppl. figures

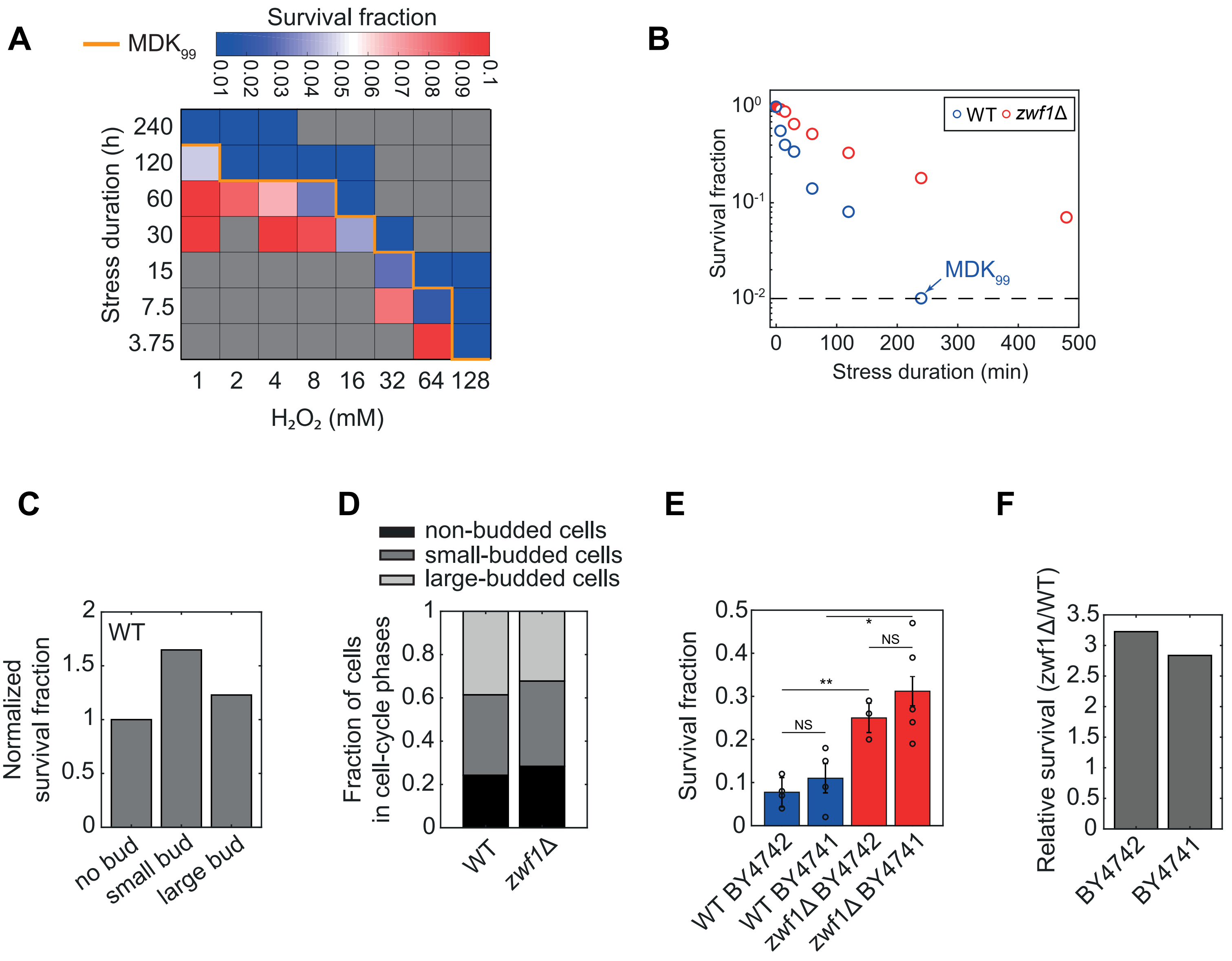
A
MDK99
Survival fraction
0.01 0.02 0.03 0.04 0.05 0.06 0.07 0.08 0.09 0.1
Stress duration (h)
240
120
60
30
15
7.5
3.75
1 2 4 8 16 32 64 128
H2O2 (mM)
B
Survival fraction
10^0
10^-1
10^-2
WT zwf1Δ
MDK99
0 100 200 300 400 500
Stress duration (min)
C
WT
Normalized survival fraction
2
1.5
1
0.5
0
no bud
small bud
large bud
D
non-budded cells
small-budded cells
large-budded cells
Fraction of cells in cell-cycle phases
1
0.8
0.6
0.4
0.2
0
WT
zwf1Δ
E
Survival fraction
0.5
0.4
0.3
0.2
0.1
0
NS
**
*
NS
WT BY4742
WT BY4741
zwf1Δ BY4742
zwf1Δ BY4741
F
Relative survival (zwf1Δ/WT)
3.5
3
2.5
2
1.5
1
0.5
0
BY4742
BY4741

**Figure S1: $H_2O_2$ tolerance in WT and zwf1 mutant strains. Related to Figures 1 and 4.**

**(A)** Heatmap of the fraction of survival cells as a function of $H_2O_2$ concentration and time under stress (N≥2 for each tested condition). Gray squares are non-tested conditions. The orange line delineates the MDK99 (minimal duration to kill 99% of the cells).

**(B)** Survival fraction of the WT (blue) and *zwf1Δ* mutant (red) above the MIC at 1 mM. One experimental repeat is represented, with n>100 per condition. The dash-line indicates the MDK99.

**(C)** Survival fraction in WT cells above the MIC (averaged from conditions ranging from 1 to 64 mM $H_2O_2$) depending on the cell cycle phase before stress exposure (non-budded, small-budded, and large-budded), relative to the small-budded cells (n = 451, 691 and 717 for non-budded, small-budded and large-budded cells respectively).

**(D)** Fraction of non-budded, small, and large-budded cells before stress exposure in WT (n = 451, 69,1 and 717 respectively) and *zwf1*Δ cells (n = 281, 390 and 319 respectively).

**(E)** Survival fraction of WT and *zwf1*Δ strains in BY4742 and BY4741 backgrounds at 32 mM for 7.5 min. N = 3 to 5 experimental replicates per condition with n>100 single-cells per replicate. Bar and error bars represent mean +/- std.

**(F)** Relative survival between WT and *zwf1*Δ strains (Survival(*zwf1*Δ)/Survival(WT)) in BY4742 and BY4741 backgrounds, based on data in E.

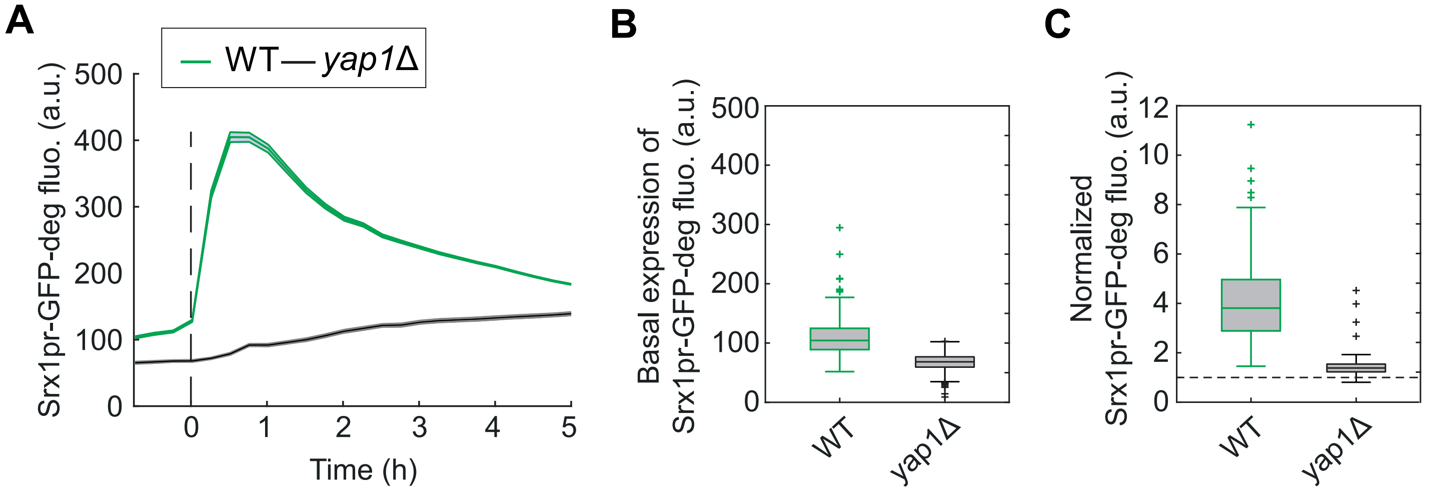
A
WT
yap1Δ
Srx1pr-GFP-deg fluo. (a.u.)
500
400
300
200
100
0
0
1
2
3
4
5
Time (h)
B
Basal expression of Srx1pr-GFP-deg fluo. (a.u.)
500
400
300
200
100
0
WT
yap1Δ
C
Normalized Srx1pr-GFP-deg fluo. (a.u.)
12
10
8
6
4
2
0
WT
yap1Δ

**Figure S2: Yap1-dependent regulation of the Srx1pr-GFP-deg expression. Related to Figure 2.**

**(A)** Mean Srx1pr-GFP-deg fluorescence +/- s.e.m. in WT and *yap1*Δ strains in response to 0.1mM $H_2O_2$ (n>100 single-cells for each strain).

**(B)** Boxplot of the basal expression (before stress exposure) of Srx1-GFP-deg in WT (n=337) and *yap1*Δ (n=118) strains.

**(C)** Upregulation of Srx1pr-GFP-degron upon a 1h stress exposure. Boxplots indicate the normalized fluorescence at t=1h over the basal fluorescence (t=0h) for WT (n=334) and *yap1*Δ (n=114) cells.

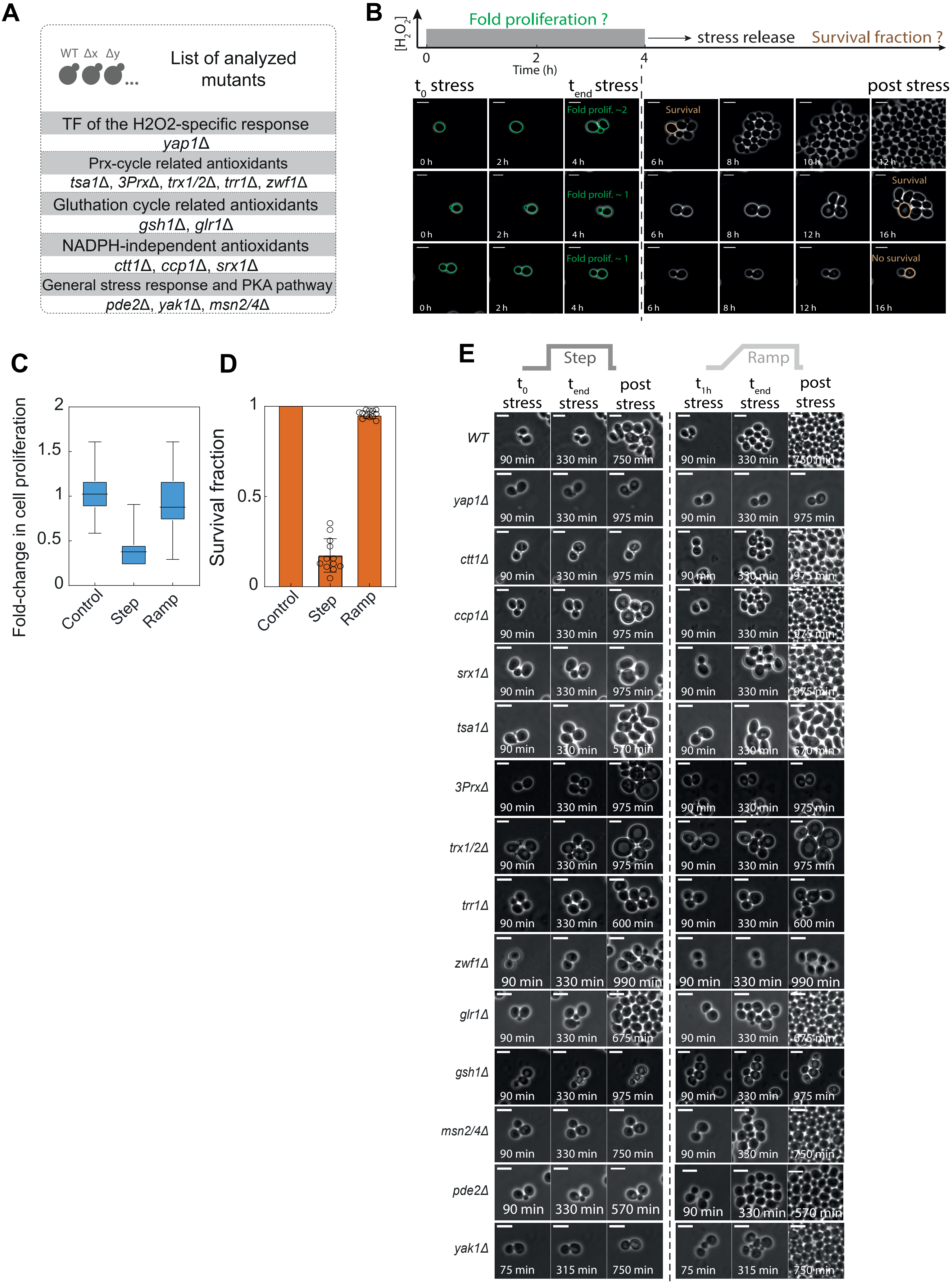
A
WT Δx Δy
List of analyzed mutants
TF of the H2O2-specific response
yap1Δ
Prx-cycle related antioxidants
tsa1Δ, 3PrxΔ, trx1/2Δ, trr1Δ, zwf1Δ
Gluthation cycle related antioxidants
gsh1Δ, glr1Δ
NADPH-independent antioxidants
ctt1Δ, ccp1Δ, srx1Δ
General stress response and PKA pathway
pde2Δ, yak1Δ, msn2/4Δ
B
[H2O2]
Fold proliferation ?
stress release
Survival fraction ?
Time (h)
t0 stress
tend stress
post stress
Fold prolif. ~2
Survival
Fold prolif. ~ 1
Survival
Fold prolif. ~ 1
No survival
C
Fold-change in cell proliferation
Control
Step
Ramp
D
Survival fraction
Control
Step
Ramp
E
Step
Ramp
t0 stress
tend stress
post stress
t1h stress
WT
yap1Δ
ctt1Δ
ccp1Δ
srx1Δ
tsa1Δ
3PrxΔ
trx1/2Δ
trr1Δ
zwf1Δ
glr1Δ
gsh1Δ
msn2/4Δ
pde2Δ
yak1Δ

**Figure S3: Resistance and tolerance in WT and stress-response mutant strains in response to step and ramp of $H_2O_2$. Related to Figure 3.**

**(A)** List of the mutant strains analyzed in this study.

**(B)** Sequences of phase contrast images of representative WT resistant (top), tolerant (middle), and dead (bottom) cells;

**(C)** Fold-change in cell proliferation under stress (based on n=55 micro-colonies, see Methods for details) for steps and ramps in WT cells.

**(D)** Post-stress survival for steps and ramps in WT cells (N = 12 independent technical replicates, n>100 for each replicate).

**(E)** Sequence of images showing representative cells of the indicated strain background in response to step/ramp of $H_2O_2$. Scale bar: 6.2 μm.

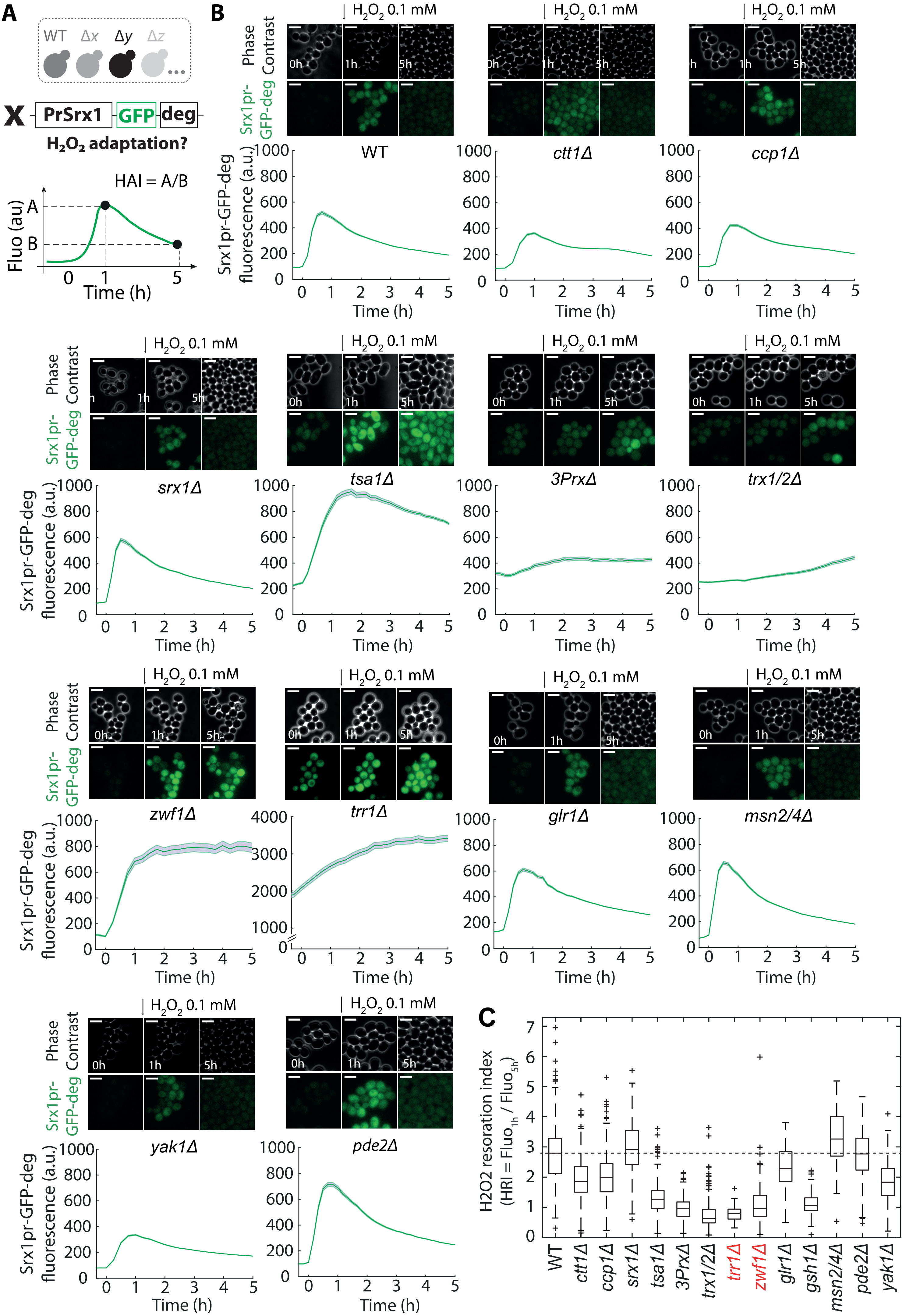
A
B
C
WT
Δx
Δy
Δz
PrSrx1
GFP
deg
H2O2 adaptation?
HAI = A/B
Fluo (au)
Time (h)
H2O2 0.1 mM
Phase Contrast
Srx1pr-GFP-deg
Srx1pr-GFP-deg fluorescence (a.u.)
WT
ctt1Δ
ccp1Δ
srx1Δ
tsa1Δ
3PrxΔ
trx1/2Δ
zwf1Δ
trr1Δ
glr1Δ
msn2/4Δ
yak1Δ
pde2Δ
H2O2 resoration index
(HRI = Fluo1h / Fluo5h)
gsh1Δ

**Figure S4: Srx1pr-GFP-deg expression in WT and stress-response mutant strains in response to $H_2O_2$. Related to Figure 3.**

**(A)** Sketch of the quantification of the $H_2O_2$ restoration index (HRI), calculated as the fluorescence 5h after stress exposure divided by the fluorescence 1h after stress exposure.

**(B)** Sequence of phase contrast and Srx1pr-GFP-degron fluorescence images (top) and quantification of the mean Srx1pr-GFP-deg expression +/- s.e.m. of a population of cells with the indicated background during a continuous 0.1 mM $H_2O_2$ exposure;

**(C)** Boxplot of the $H_2O_2$ restoration index (HRI = Fluo(1h) / Fluo(5h)) for the indicated mutant strains. The two mutants with the trade-off between $H_2O_2$ resistance and tolerance (*trr1*Δ and *zwf1*Δ) are represented in red.

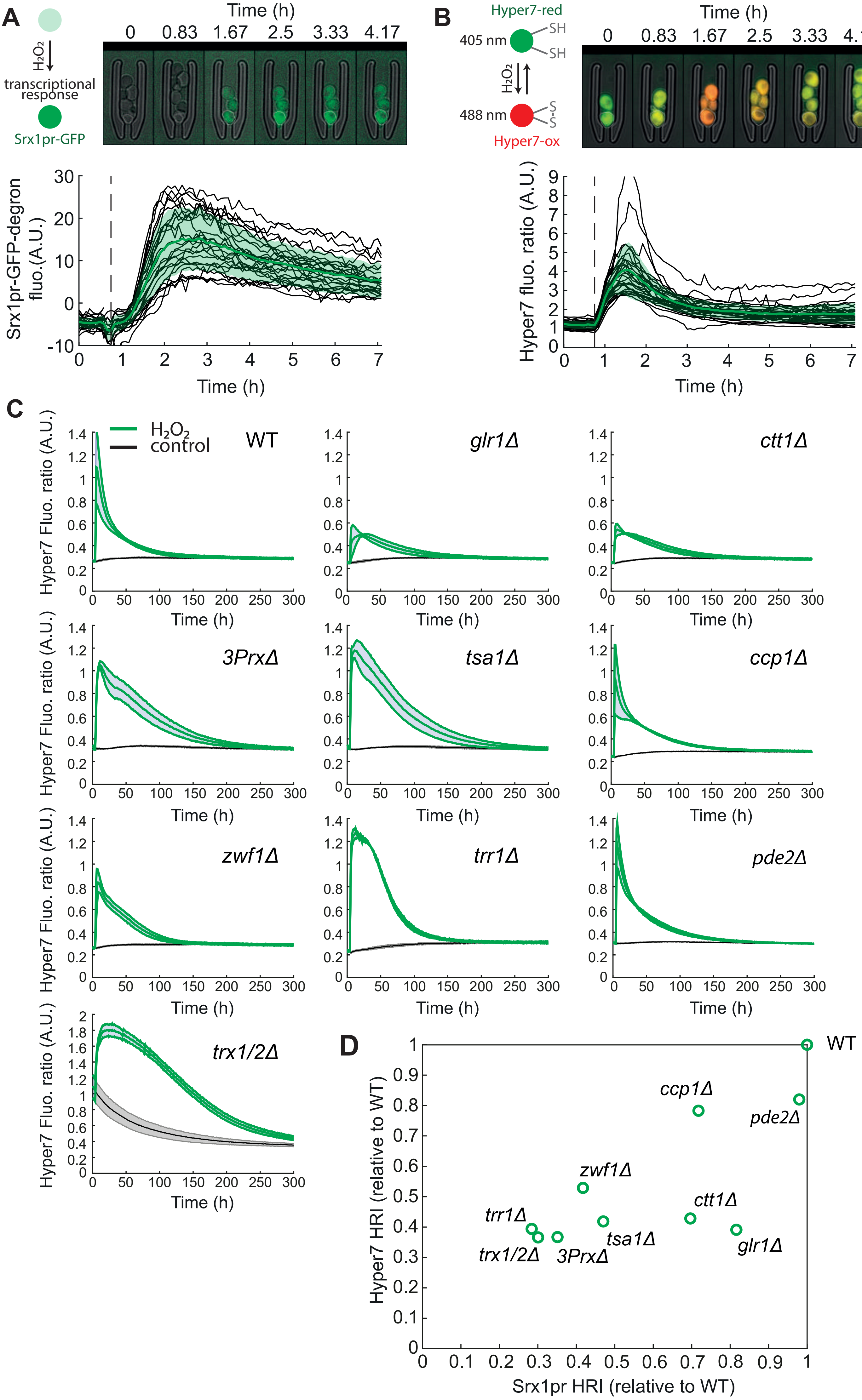
A
$H_2O_2$
transcriptional response
Srx1pr-GFP
Time (h)
0 0.83 1.67 2.5 3.33 4.17
Srx1pr-GFP-degron fluo.(A.U.)
Time (h)
B
Hyper7-red
405 nm
SH
SH
$H_2O_2$
488 nm
S-S
Hyper7-ox
Hyper7 fluo. ratio (A.U.)
C
$H_2O_2$
control
WT
glr1Δ
ctt1Δ
3PrxΔ
tsa1Δ
ccp1Δ
zwf1Δ
trr1Δ
pde2Δ
trx1/2Δ
Hyper7 Fluo. ratio (A.U.)
Time (h)
D
Hyper7 HRI (relative to WT)
Srx1pr HRI (relative to WT)

**Figure S5: Hyper7 probe expression in WT and mutants and comparison to the Srx1pr-GFP-deg expression. Related to Figure 3.**

**(A)** Top panel: schematic of the Srx1pr-GFP-degron expression, with a time series of WT cells expressing Srx1pr-GFP-deg in the microfluidic chip. Bottom panel: Quantification of the Srx1pr-GFP-deg expression over time upon $H_2O_2$ stress exposure. Single-cell trajectories are shown in black, with the mean fluorescence expression ± standard deviation (n = 30 cells) represented in green. The dashed black line marks the onset of the 0.4 mM $H_2O_2$ exposure.

**(B)** Top panel: Schematic representation of the Hyper7 probe, accompanied by a time series of WT cells expressing Hyper7 in the microfluidic chip. Bottom panel: Quantification of the Hyper7 ratio expression (488 nm/405 nm; see Methods) over time following $H_2O_2$ stress exposure. Single-cell trajectories are shown in black, while the mean fluorescence expression ± standard deviation (n = 30 cells) is represented in green. The dashed black line marks the onset of the 0.4 mM $H_2O_2$ exposure.

**(C)** Expression of the Hyper7 probe in the different mutants under 0.2 mM (blue curve +/- standard deviation of the mean of N=3 experimental replicates) or without $H_2O_2$ (black curve +/- standard deviation of the mean of N=3 experimental replicates).

**(D)** Comparison of the HRI values derived from the Srx1pr-GFP-degron signal and the Hyper7 probe across nine mutants from the screen shown in Figure 3, normalized relative to the WT for each probe.

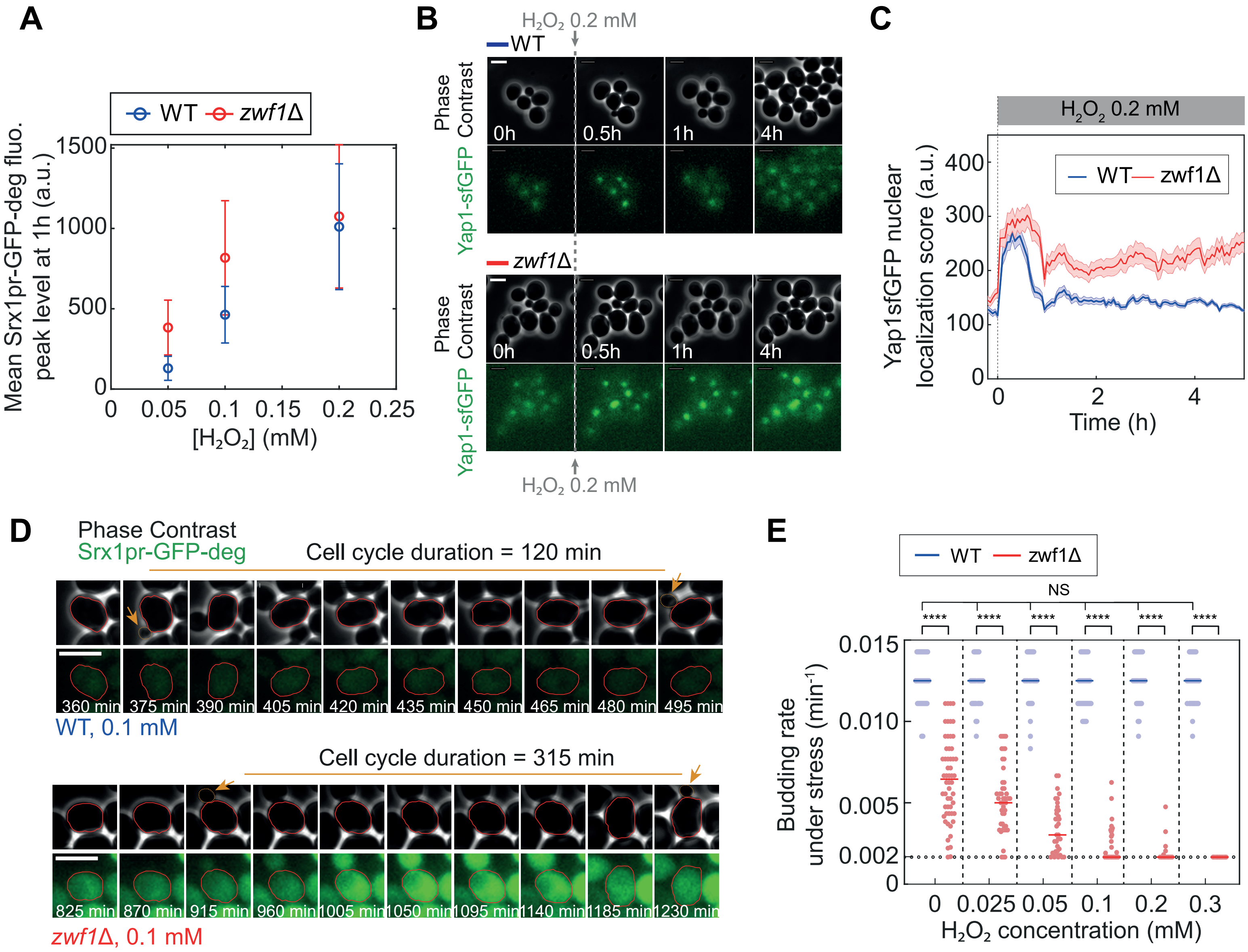
A
WT zwf1Δ
Mean Srx1pr-GFP-deg fluo. peak level at 1h (a.u.)
[H2O2] (mM)
B
H2O2 0.2 mM
WT
Phase Contrast
Yap1-sfGFP
0h
0.5h
1h
4h
zwf1Δ
C
H2O2 0.2 mM
Yap1sfGFP nuclear localization score (a.u.)
Time (h)
D
Phase Contrast
Srx1pr-GFP-deg
Cell cycle duration = 120 min
WT, 0.1 mM
Cell cycle duration = 315 min
zwf1Δ, 0.1 mM
E
Budding rate under stress (min-1)
H2O2 concentration (mM)
NS
****

**Figure S6: Relation between $H_2O_2$ homeostasis and cell growth in WT and *zwf1Δ* strains. Related to Figure 4.**

**(A)** Mean Srx1pr-GFP-deg fluorescence +/- std measured 1 h after the onset of stress exposure in WT and *zwf1Δ* strains as a function of $H_2O_2$ concentration.

**(B)** Sequence of phase contrast and Yap1-sfGFP fluorescence in WT and *zwf1Δ* strains in response to a 0.2 mM $H_2O_2$ exposure.

**(C)** Yap1-sfGFP nuclear localization score +/- s.e.m. in WT and *zwf1Δ* cells measured from the experiment in (B). The vertical dashed line indicates the onset of stress exposure.

**(D)** Sequence of phase contrast and Srx1pr-GFP-deg fluorescence of WT and zwf1Δ cells continuously exposed to a 0.1 mM $H_2O_2$ stress. Orange arrows indicate budding events.

**(E)** Budding rates of WT (blue symbols) and *zwf1Δ* (red symbols) cells as a function of $H_2O_2$ concentration; each filled symbol corresponds to a single cell; horizontal colored lines represent the median budding rate. All the cell cycles with a frequency smaller than 0.0017 $min^{-1}$ were included in the analysis as of a frequency of 0.0017 $min^{-1}$. Statistical differences (two-sided Mann-Whitney U test) are indicated by **** for $p < 0.0001$, NS for $p > 0.05$.

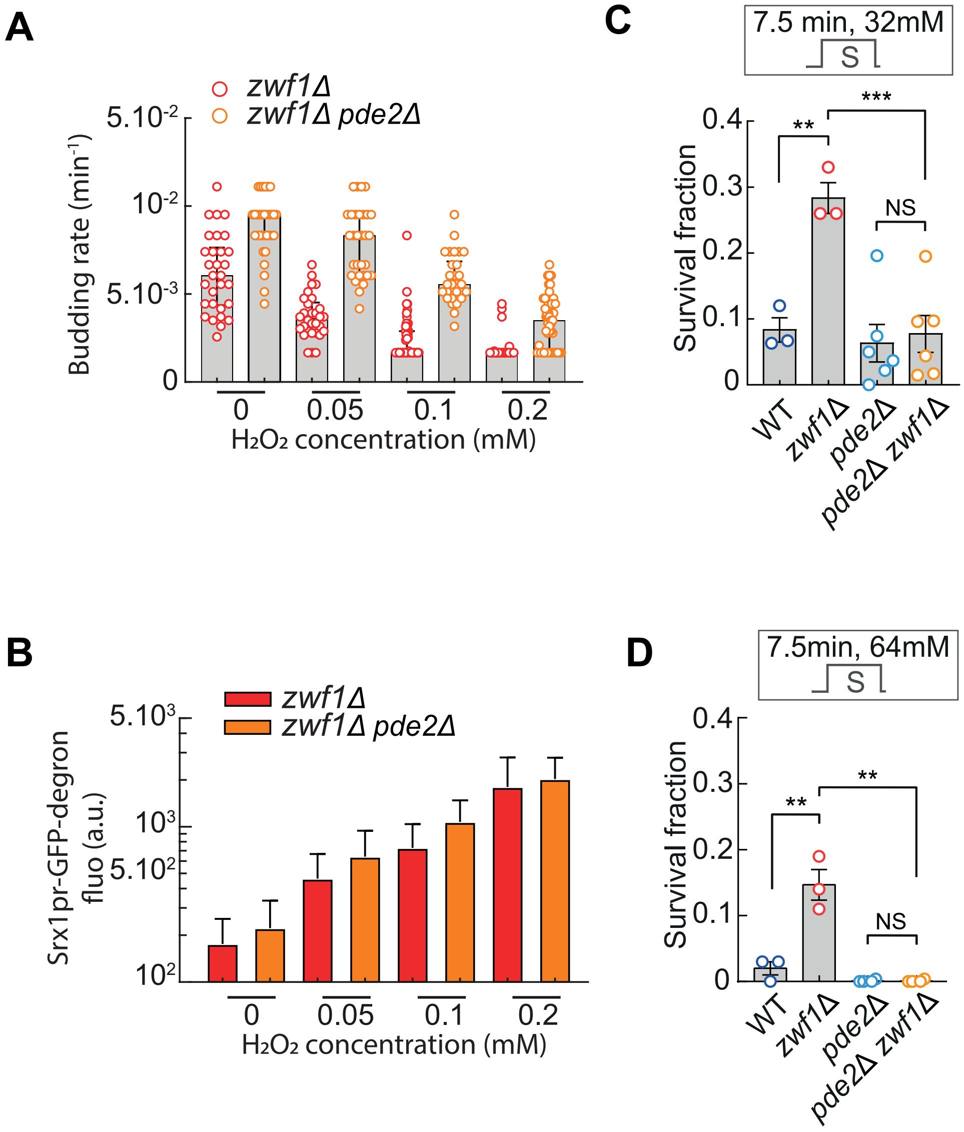

A
zwf1Δ
zwf1Δ pde2Δ
Budding rate (min-1)
5.10-2
10-2
5.10-3
0
0
0.05
0.1
0.2
H2O2 concentration (mM)
C
7.5 min, 32mM
S
Survival fraction
0.4
0.3
0.2
0.1
0
**
***
NS
WT
zwf1Δ
pde2Δ
pde2Δ zwf1Δ
B
zwf1Δ
zwf1Δ pde2Δ
Srx1pr-GFP-degron fluo (a.u.)
5.103
103
5.102
102
0
0.05
0.1
0.2
H2O2 concentration (mM)
D
7.5min, 64mM
S
Survival fraction
0.4
0.3
0.2
0.1
0
**
**
NS
WT
zwf1Δ
pde2Δ
pde2Δ zwf1Δ

**Figure S7: The PKA pathway modulates $H_2O_2$ tolerance and resistance in a Prx-dependent manner. Related to Figure 5.**

**(A)** Budding rate of individual cells (*zwf1Δ*, open red circles; *zwf1Δ pde2Δ*, open orange circles) under increasing $H_2O_2$ concentrations. Budding rates are normalized as shown in Figure 5F. The sample size ranges from 31 to 105 single cells.

**(B)** Mean Srx1pr-GFP-degron expression ± s.e.m. in the *zwf1Δ* (red bars) and *zwf1Δ pde2Δ* (orange bars) strains under corresponding $H_2O_2$ concentrations (log scale), measured 5 hours after stress onset. Sample sizes are n > 242 for *zwf1Δ* and n > 246 for *zwf1Δ pde2Δ* across all concentrations.

**(C)** and **(D)** Survival fraction in response to $H_2O_2$ boluses (32 or 64 mM for 7.5 min). Filled circles represent independent technical replicates (N = 3 to 6, n>100 for each replicate), error bars are s.e.m. Statistical analysis is based on a one-sided t-test for comparison between WT and *zwf1*Δ and between *pde2*Δ and *zwf1*Δ*pde2*Δ, and a two-sided t-test for comparison between *zwf1*Δ and *zwf1*Δ*pde2*Δ (no assumption was made on the effect of *PDE2* deletion in the *zwf1*Δ background).

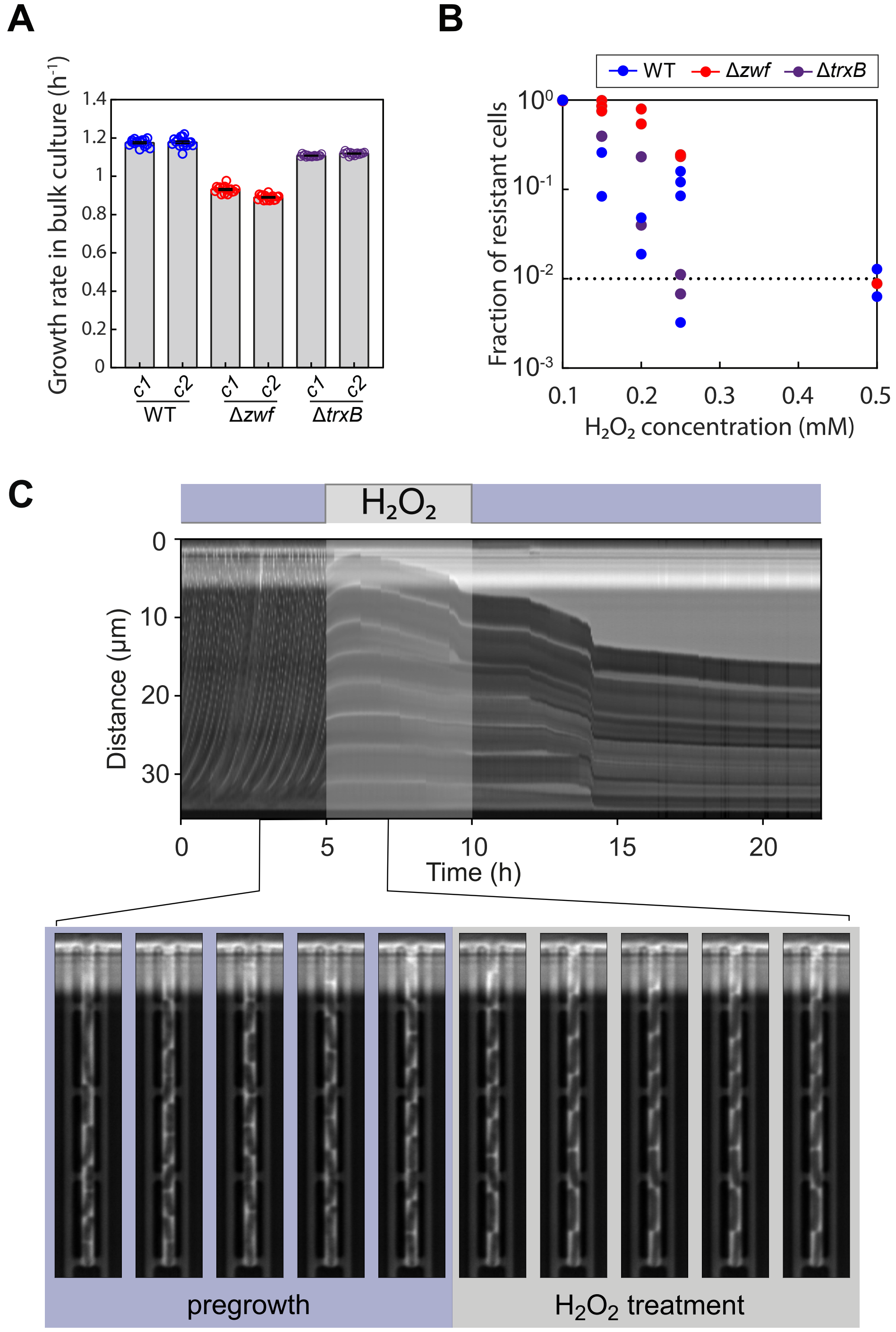

A
B
C
Growth rate in bulk culture (h$^{-1}$)
1.4
1.2
1
0.8
0.6
0.4
0.2
0
c1
c2
WT
Δ*zwf*
Δ*trxB*
WT
Δ*zwf*
Δ*trxB*
Fraction of resistant cells
$10^0$
$10^{-1}$
$10^{-2}$
$10^{-3}$
0.1
0.2
0.3
0.4
0.5
$H_2O_2$ concentration (mM)
$H_2O_2$
Distance (µm)
0
10
20
30
0
5
10
15
20
Time (h)
pregrowth
$H_2O_2$ treatment

**Figure S8: Conservation of the NADPH-dependent trade-off between H2O2 stress resistance and tolerance in the prokaryote *E. coli*. Related to Figure 6.**

**(A)** Growth rate of WT, Δ*zwf,* and Δ*trxB* strains in liquid cultures. c1 and c2 are two different clones. Circles show the different technical replicates and the gray bars indicate the mean over the replicates.

**(B)** Sequence of raw phase contrast images of *E. coli* cells in the microfluidic device (bottom) and reduction of the dimensionality using a kymograph (top image).

**(C)** Fraction of resistant cells in WT, Δ*zwf* and Δ*trxB* strains. Circles represent technical replicates (N = 2 to 4 for concentrations close to the MIC). The dashed line indicates a 1% survival, from which the MIC can be calculated.
